# Exploiting the SunTag system to study the developmental regulation of mRNA translation

**DOI:** 10.1101/2025.02.09.637317

**Authors:** Alastair Pizzey, Catherine Sutcliffe, Jennifer C. Love, Emmanuel Akabuogu, Magnus Rattray, Mark P. Ashe, Hilary L. Ashe

## Abstract

The ability to quantitatively study mRNA translation using SunTag imaging is transforming our understanding of the translation process. Here, we expand the SunTag method to study new aspects of translation regulation in *Drosophila*. Repression of the maternal *hunchback* (*hb*) mRNA in the posterior of the *Drosophila* embryo is a textbook example of translational control. Using SunTag imaging to quantitate translation of maternal *SunTag-hb* mRNAs, we show that repression in the posterior is leaky as ∼5% of *SunTag-hb* mRNAs are translated. In the anterior of the embryo, the maternal and zygotic *SunTag-hb* mRNAs show similar translation efficiency despite having different UTRs. We demonstrate that the *SunTag-hb* mRNA can be used as a reporter to study ribosome pausing at single-mRNA resolution, by exploiting the conserved *xbp1* mRNA and A60 pausing sequences. Finally, we adapt the detector component of the SunTag system to visualise and quantitate translation of the *short gastrulation* (*sog*) mRNA, encoding an essential secreted extracellular BMP regulator, at the endoplasmic reticulum in fixed and live embryos. Together, these tools will facilitate the future dissection of translation regulatory mechanisms during development.

## Introduction

Translational regulation is critical for development and homeostasis (Kong and Lasko, 2012; Lasko, 2020; Tahmasebi et al., 2019). There are three main steps to cap-dependent translation: initiation, elongation and termination (Brito Querido et al., 2024; Hellen, 2018; Knight et al., 2020; Merrick and Pavitt, 2018). Initiation of cap-dependent translation involves assembly of the pre-initiation complex, which consists of the small 40S ribosome subunit, the initiator methionyl-tRNA and a number of initiation factors. This complex is recruited to the 5’ end of the mRNA via interaction with mRNA bound translation factors, such as eukaryotic initiation factor (eIF)4E and eIF4G. The pre-initiation complex then scans along the mRNA until the start codon is reached and the large 60S subunit joins to form the translationally competent ribosome (Brito Querido et al., 2024; Merrick and Pavitt, 2018). Elongation is the repeated process in which elongation factors bring aminoacyl-tRNAs to the ribosome, the amino acid is added to the growing nascent polypeptide chain and the ribosome moves to the next codon (Knight et al., 2020). Termination occurs when the stop codon is recognised by termination factors which lead to release of the synthesised protein and disassembly of the ribosome (Hellen, 2018).

Translation initiation, elongation and termination are all tightly controlled (Brito Querido et al., 2024; Hellen, 2018; Knight et al., 2020; Merrick and Pavitt, 2018) and misregulation of these processes is associated with human diseases, including cancer and neurodevelopment disorders (Jishi et al., 2021). Of the three steps, the control of initiation has been the most highly studied. Many RNA binding proteins and microRNAs have been implicated in the control of translation initiation (Harvey et al., 2018; Wilczynska and Bushell, 2015). Examples of RNA binding proteins that repress translation initiation are eIF4E homologous protein (4EHP) and eIF4E binding proteins. 4EHP binds to the mRNA 5’ cap but, unlike eIF4E, is unable to interact with eIF4G so translation initiation is repressed (Christie and Igreja, 2023). eIF4E binding proteins, such as *Drosophila* Cup, inhibit translation initiation by binding to eIF4E and preventing its interaction with eIF4G (Lasko, 2020).

Although less intensively studied, translation elongation is also an important regulated step (Knight et al., 2020). There is considerable variation in the rate at which mRNA codons are decoded by the ribosome, with more time needed for the ribosome to decode particular codons or regions of the mRNA at ribosomal pauses. Ribosome pausing can be caused by rare codons, RNA structures, RNA binding proteins, mRNA modifications, tRNA/amino acid deficiency and nascent protein interactions in the ribosome exit tunnel (Buskirk and Green, 2017). These delays in ribosome movement can have important physiological roles, for example by facilitating protein targeting or folding of the nascent polypeptide (Gloge et al., 2014). In addition, paused ribosomes and the resulting collided ribosomes can act as a diagnostic signature for the cell to identify problems such as RNA damage or aberrant nascent protein folding, which ultimately target the mRNA and nascent protein chain for degradation (Inada, 2020).

A classical ribosome pause site is found in the *xbp1* mRNA. In unstressed mammalian cells, the ribosome pauses at a specific site during translation of the unspliced *XBP1* mRNA (Yanagitani et al., 2011). The nascent polypeptide that has been translated (XBP1u) targets the mRNA to the endoplasmic reticulum (ER) membrane. Following ER stress, the IRE1α transmembrane protein is recruited, which promotes cytoplasmic splicing of the *XBP1* mRNA. This splicing causes a frameshift in the *XBP1s* mRNA so that the XBP1s protein is translated with a distinct C-terminal half of the protein (Yanagitani et al., 2009). XBP1s is a transcription factor that induces transcription of genes involved in alleviating ER stress (Yoshida et al., 2001). The *xbp1* mRNA pause site is broadly conserved in organisms ranging from yeast to mammals (Chyżyńska et al., 2021).

Another example of ribosomal pausing is induced by stretches of adenosines, which encode multiple lysine residues. This pausing is due to interactions between the poly(A) mRNA and nascent poly-Lys stretch with the ribosome (Chandrasekaran et al., 2019; Koutmou et al., 2015; Tesina et al., 2020). Poly(A) sequences have been used to induce ribosome collisions, in order to study the quality control of translation in mammalian cells (Goldman et al., 2021).

The SunTag method is a powerful technique that allows translation to be studied in fixed and live cells at single mRNA resolution (Pichon et al., 2016; Tanenbaum et al., 2014; Wang et al., 2016; Wu et al., 2016; Yan et al., 2016). Briefly, this method uses a single chain variable fragment (scFv) fused to a fluorescent protein (FP) to detect an array of SunTag peptides inserted at the N-terminus of the protein of interest (Fig. 1Ai). The mRNA can be co-detected either using single molecule fluorescent in situ hybridisation (smFISH) in fixed cells or the MS2 system in living cells.

**Fig 1.**
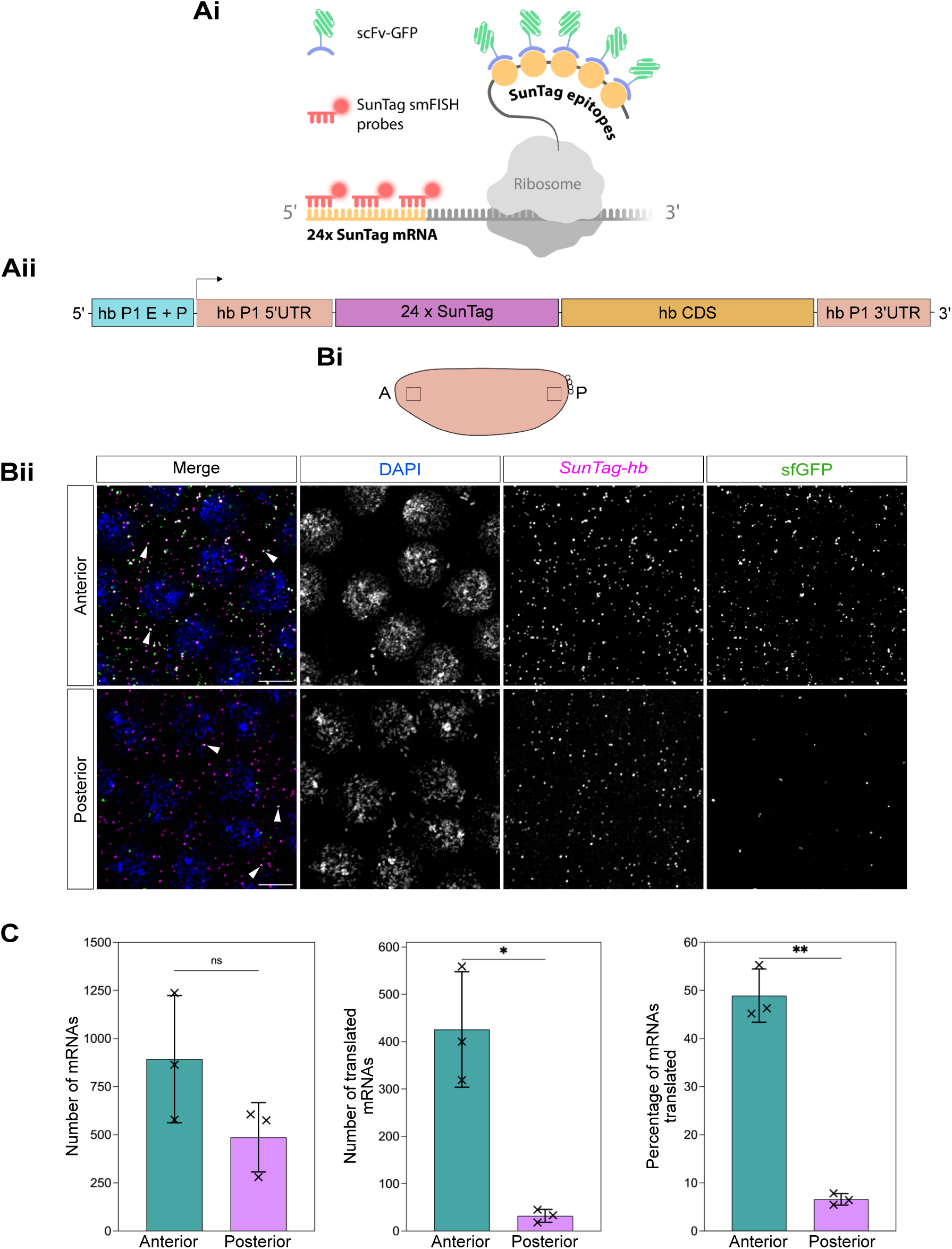
Leaky repression of maternal *24xSunTag-hb* mRNA translation in the posterior of the embryo. (A) (i) Schematic showing *24xSunTag-hb* mRNA translation. Translated SunTag epitopes are recognised by a scFv-sfGFP fusion protein, mRNAs are detected using SunTag smFISH probes. (ii) Schematic of the maternal *matE>24xSunTag*-*hb* transgene. hb P1 E + P = *hb* maternal P1 enhancer and promoter. (B) (i) Schematic shows the positions imaged in the embryo anterior and posterior, with the ubiquitous distribution of maternal *hb* mRNAs in peach. (ii) Representative high magnification images from a region in the anterior and posterior of a nc13 embryo from *matα4-GAL4-VP16 UASp-scFv-sfGFP-NLS/matE>24xSunTag-hb* females showing scFv-sfGFP signal (green) and stained with DAPI (blue) and *SunTag* smFISH probes (magenta). White arrowheads show translation sites based on colocalised *24xSunTag-hb* and scFv-sfGFP signals. Scale bars: 5 μm. (C) Quantitation of the number of cytoplasmic mRNAs, translated mRNAs and percentage of translated mRNAs in anterior (cyan) and posterior (magenta) regions of interest in nc13 embryos, for 3 biological repeats (including the images shown in (Bii)). Means ± SD, Paired two-tailed Student’s *t-*test, ns - not significant, \**P*<0.05, \*\**P*<0.01.

The SunTag system has been used to study different aspects of mRNA translation in *Drosophila*. In the *Drosophila* early embryo, zygotic *SunTag-hb* (Vinter et al., 2021a) and *SunTag-nanos* (Chen et al., 2024) mRNAs were found to adopt more open conformations with a greater separation between their 5’ and 3’ ends when they are being translated, compared to the untranslated mRNAs. Studies of translation of the *twist* and *Insulin-like peptide 4* mRNAs in the *Drosophila* embryo by the SunTag method revealed spatial heterogeneities in their translation efficiency depending on the apical-basal position in the cell (Dufourt et al., 2021). Imaging of *SunTag-nanos* mRNAs demonstrated that they are translated in germ cell granules in the *Drosophila* embryo, where Oskar sequesters the Smaug translational repressor that prevents non-localised *SunTag-nanos* mRNAs from being translated in the soma (Chen et al., 2024). Finally, the neuromodulator Tyramine leads to decondensation of RNP granules in *Drosophila* brain mushroom body neurons, resulting in the translation activation of granule-associated *SunTag-profilin* mRNAs (Formicola et al., 2021).

Despite the progress made by studying translation during development using the SunTag system, there are many other areas of translational control that have yet to be explored. Here we show the versatility of the system by providing proof-of-principle data for new avenues of exploration. These include quantitation of translation repression strength, comparisons of translation efficiencies of maternal vs zygotic transcripts for the mRNA of interest, studies of ribosome pausing and visualisation of translation of mRNAs encoding secreted proteins.

## Results

### Incomplete maternal *hb* mRNA translation repression in the posterior of the embryo

The Hb transcription factor is critical for anterior-posterior patterning (Tautz et al., 1987). While we studied translation of zygotic *24xSunTag-hb* mRNAs in our previous study (Vinter et al., 2021a), *hb* is also maternally expressed (Bender et al., 1988; Tautz et al., 1987). *hb* mRNAs are maternally deposited throughout the embryo but translationally repressed in the posterior by RNA binding proteins (Hülskamp et al., 1989; Hülskamp et al., 1990; Irish et al., 1989; Murata and Wharton, 1995; Sonoda and Wharton, 2001; Struhl et al., 1989; Tautz, 1988; Wharton and Struhl, 1991; Zamore et al., 1997). To study translation of maternal *hb* mRNAs, we generated a transgene in which *24xSunTag-hb* coding sequences are transcribed under the control of the maternal P1 enhancer and promoter, with the maternal mRNA UTRs (Lukowitz et al., 1994; Margolis et al., 1995; Schröder et al., 1988) (Fig. 1Aii). In the experiments presented here, we use the *matα4-GAL4-VP16* driver to induce high level maternal expression of a *UASp-scFv-sfGFP-NLS* transgene. Embryos were collected from females carrying the maternal *matE>24xSunTag-hb* transgene, plus *matα4-GAL4-VP16* and *UASp-scFv-sfGFP-NLS* insertions. Maternal *24xSunTag-hb* mRNAs were detected by smFISH and translation sites were visualised based on the scFv-sfGFP signal in fixed embryos (Fig. 1Ai).

To determine the strength of translational repression of maternal *24xSunTag-hb* mRNAs in the posterior of the embryo at nc13, we imaged a region of interest in the anterior and posterior of the embryo (Fig. 1Bi, ii). Individual mRNAs and translation sites were quantitated, with translation sites identified based on the colocalisation of scFv-sfGFP fluorescent signals with mRNAs. Quantitation of the total number of mRNAs in each region showed a trend of lower mRNA numbers in the posterior, although the difference is not significant (Fig. 1C). In the anterior region of the embryo, many translation sites are evident, with ∼50% of the maternal *24xSunTag-hb* mRNAs translated (Fig. 1C). There is a significant reduction in the proportion of maternal *24xSunTag-hb* mRNAs translated in the posterior, where only a small number of weak translation sites are detected (Fig. 1C).

We also used a GCN4 antibody, which recognises the SunTag epitopes, and SunTag smFISH probes to detect translation sites in fixed embryos carrying one copy of the maternal *24xSunTag-hb* transgene (Fig. S1A). These *matE>24xSunTag-hb* heterozygous embryos lacked the *scFv-sfGFP-NLS* transgene, to avoid scFv-sfGFP proteins from blocking anti-GCN4 from binding to the SunTag epitopes. In maternal *24xSunTag-hb* nc13 embryos, the proportion of mRNAs translated in the anterior is similar (∼60%) to that observed using scFv-sfGFP detection (Fig. S1B). In the posterior of the embryo, ∼3% of maternal *24xSunTag-hb* mRNAs are translated (Fig. S1B), again similar to the proportion translated (∼6%) calculated based on scFv-sfGFP detection (Fig. 1C). In addition, the number of maternal *24xSunTag-hb* mRNAs present in the posterior is significantly lower than in the anterior (Fig. S1B), suggesting that translation repression may lead to mRNA degradation.

It is possible that a small proportion (∼3-6%) of maternal *24xSunTag-hb* mRNAs are translated in the embryo posterior because the increased number of *24xSunTag-hb* mRNAs, which contain the endogenous *hb* UTRs, titrates out a repressor that is otherwise present at sufficient concentration to completely repress endogenous *hb* mRNAs. To test this, we used the GCN4 antibody and SunTag smFISH probes to image translation in embryos homozygous for the maternal *SunTag-hb* transgene, which have a further increase in the number of *24xSunTag-hb* mRNAs (Fig. S1C, D). Quantitation of the data shows that the proportion of maternal *24xSunTag-hb* mRNAs translated in the anterior and posterior regions of homozygous embryos is similar to that detected in heterozygous embryos (Fig. S1D, cf Fig. S1B), rather than being dramatically increased as predicted if a translational repressor was limiting. In addition, as observed for the heterozygous *matE>24xSunTag-hb* embryos (Fig. S1B), there are significantly fewer maternal *24xSunTag-hb* mRNAs in the posterior region compared to the anterior (Fig. S1D). Together these results show that translation repression of maternal *24xSunTag-hb* mRNAs is not absolute in the posterior of the embryo and suggest that this is unlikely to be due to titration of a translational repressor by the transgenic copies of the maternal transgene.

### Translation efficiencies of the *hb* maternal vs zygotic mRNAs

The maternal and zygotic *hb* mRNAs have the same coding sequence but different 5’ and 3’ UTRs (Margolis et al., 1995; Schröder et al., 1988) (Fig. 2A). The same ribosome elongation rate would be predicted based on the same coding sequence, but it was unclear whether the distinct UTRs might lead to different initiation rates (Leppek et al., 2018; Mayr, 2017), which would alter the number of ribosomes/mRNA. To address this, we visualised and quantitated the SunTag translation signals in an anterior region (Fig. 2Bi) to estimate ribosome numbers for the maternal and zygotic *24xSunTag-hb* mRNAs across early embryonic development (Fig. 2Bii). For maternal *24xSunTag-hb* mRNAs, embryos were collected from *matα4-GAL4-VP16 UASp-scFv-sfGFP-NLS* females also carrying one copy of the *matE>24xSunTag-hb* transgene. To study translation of zygotic *SunTag-hb* mRNAs, females carrying *matα4-GAL4-VP16* and *UASp-scFv-sfGFP-NLS* insertions were crossed to males homozygous for the *hbP2>24xSunTag-hb* transgene and the resulting embryos collected. To estimate ribosome number, the fluorescent signal at each translation site was divided by the mean fluorescence of a single protein, then this value was divided by the number of mRNAs in the translation site to control for two or more mRNAs being co-localised in the translation site (see Methods). In our previous analysis, we specifically focussed on the strongest translation sites (Vinter et al., 2021a), whereas here we set a lower threshold to include all translation sites with two or more ribosomes.

**Fig 2.**
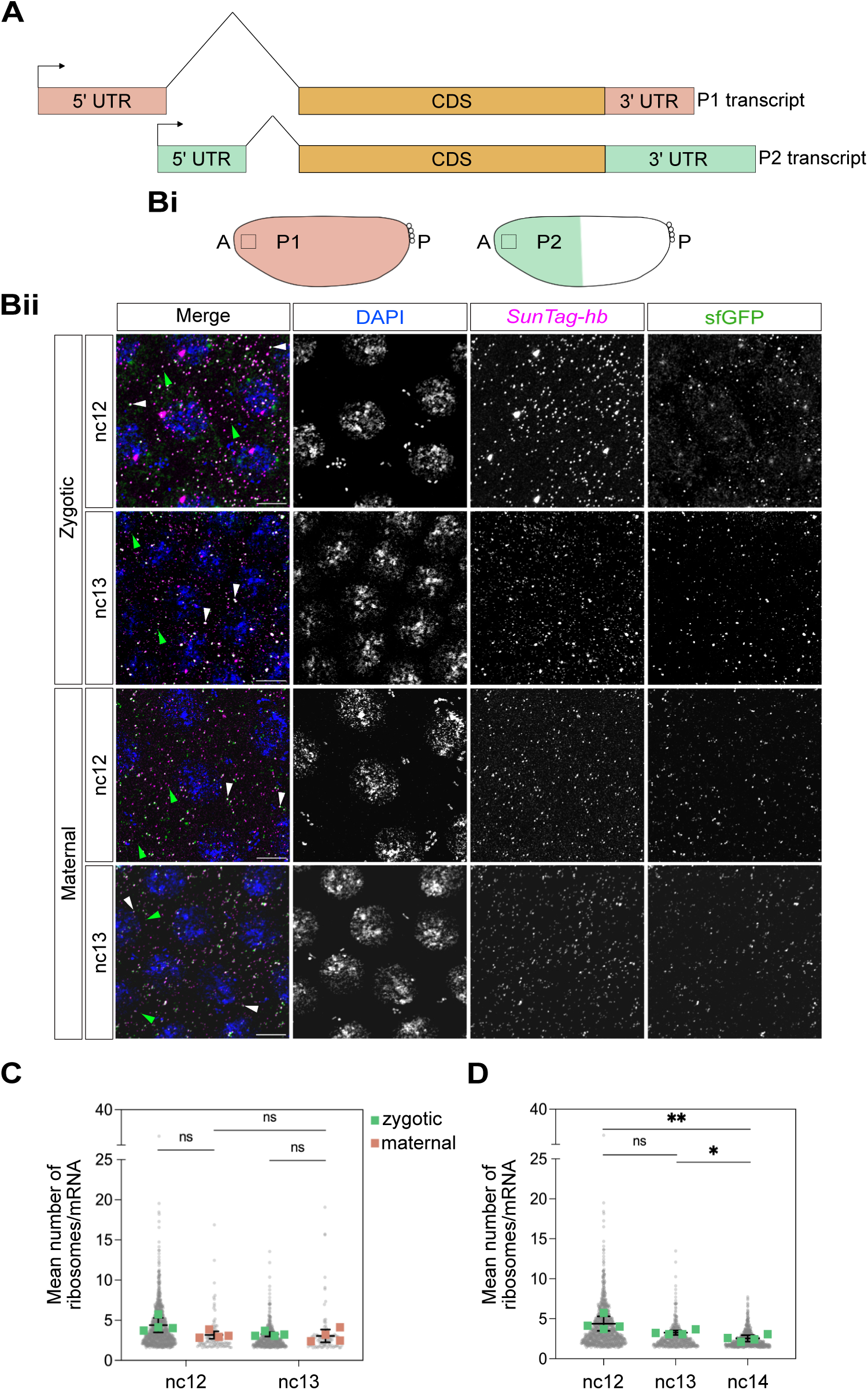
Translation efficiencies of maternal and zygotic *24xSunTag-hb* mRNAs across embryogenesis. (A) Structure of the maternal P1 and zygotic P2 *hb* transcripts. (B) (i) Schematic of embryos with the ubiquitous maternal (P1) *hb* mRNAs in peach (left) and the zygotic *hb* expression domain driven by the P2 enhancer in green (right). The position imaged in the anterior is marked with a box. (ii) High magnification images from the anterior of nc12 and nc13 embryos collected from either *matα4-GAL4-VP16 UASp-scFv-sfGFP-NLS/matE>SunTag-hb* females or *matα4-GAL4-VP16 UASp-scFv-sfGFP-NLS* females crossed to *hbP2>24xSunTag-hb* males, to detect maternal and zygotic *24xSunTag-hb* mRNAs, respectively. Merged images show detection of scFv-sfGFP signal (green), DAPI (blue) and *SunTag* smFISH probes (magenta), with single channel images for clarity. White arrowheads show colocalised scFv-sfGFP and *24xSunTag-hb* signals representing translated mRNAs. Green arrowheads show weaker non-colocalised scFv-sfGFP signals, representing single proteins. Scale bars: 5 μm. (C) Graph shows all the individual data points and mean numbers, from 3 biological replicates, of ribosomes detected on zygotic (green) and maternal (pink) *24xSunTag-hb* mRNAs in the anterior of nc12 and nc13 embryos. n=4, data are mean ± s.d. Unpaired two-tailed *t*-test, ns – not significant, \**P<*0.05. (D) Graph shows all the individual data points and mean numbers, from 3 biological replicates, of ribosomes present on zygotic *24xSunTag-hb* mRNAs in the anterior of nc12, nc13 and early nc14 embryos. The data for nc12 and nc13 are also shown in (C). n=4, data are mean ± s.d. Unpaired two-tailed *t*-test, ns – not significant, \**P<*0.05, \*\**P<*0.01.

Analysis of ribosome number at nc12 and nc13 shows that there is no significant difference in the number of ribosomes detected on zygotic *24xSunTag-hb* mRNAs compared to maternal (Fig. 2C). Similarly, there is no significant difference in ribosome number on either the maternal or zygotic *24xSunTag-hb* mRNAs between nc12 and nc13 (Fig. 2C). However, a decrease in ribosome number, indicative of a lower translation efficiency, is observed in early nc14 embryos for zygotic *24xSunTag-hb* mRNAs (Fig. 2D). We have not quantitated ribosome number for maternal *24xSunTag-hb* mRNAs at nc14 as very few mRNAs remain at this stage. Together, these data show that the maternal and zygotic *24xSunTag-hb* mRNAs have similar translation efficiencies, while that of zygotic *24xSunTag-hb* mRNAs decreases during nc14.

Our estimates of ribosome number vary over a ∼2-fold range, which appears to be dependent on how the scFv-FP protein is expressed (using the *nanos* (*nos*) enhancer and promoter vs GAL4 amplification). Analysis of images of zygotic *24xSunTag-hb* nc13 embryos expressing scFv-mNeonGreen(NG)-NLS or scFv-msGFP2-NLS under the control of the *nos* promoter gives estimates of ∼8 ribosomes/mRNA, whether anti-NG staining (of scFv-NG bound to SunTag) or fluorescence intensity is used to estimate the signal at the translation sites (Fig. S2). This contrasts with lower estimates of ∼3-4 ribosomes/mRNA from images of nc13 zygotic *24xSunTag-hb* embryos expressing *UASp-scFv-sfGFP-NLS* or *UASp-scFv-msGFP2-NLS* transgenes using the *matα4-GAL4-VP16* or *nos-GAL4-VP16* drivers, respectively (Fig. 2D, Fig. S2). As expression of scFv-msGFP2 gives different ribosome number estimates, dependent on whether scFv-msGFP2 is expressed using the *nos* promoter or GAL4 amplification, we conclude that the reason for the difference is not fluorescent protein dependent but instead the mode of expression. We speculate that the lower ribosome number estimates obtained with GAL4 amplification, despite much higher scFv-msGFP2 protein (Bellec et al., 2024), may relate to reduced signal to noise due to the higher background. Based on these observations, we prefer to use ribosome number estimates to observe relative differences, rather than make conclusions based on absolute numbers.

### Zygotic *hb* mRNA translation

During the course of this analysis, we used the zygotic *24xSunTag-hb* transgene (Fig. 3A) to quantitate the proportion of *24xSunTag-hb* mRNAs translated in early embryos from *matα4-GAL4-VP16 UASp-scFv-sfGFP-NLS* females. mRNAs and translation sites were assigned to the closest nucleus allowing us to calculate the total number of mRNAs and translation sites for a nuclear territory. Each nuclear territory is equivalent to a virtual cell, as the embryo only starts to cellularise in early nc14. Analysis of the data reveals uniform translation of zygotic *SunTag-hb* mRNAs over most of the expression domain in early nc14 embryos (Fig. S3A-C). We report the data as images (Fig. S3A) and heatmaps (Fig. S3B) for a representative embryo and as graphs (Fig. S3C) showing the mean values, based on data from 3 biological repeats, across the AP embryo axis. In contrast to the uniform translation detected here in early nc14 embryos (Fig. S3A-C), we had previously detected only a stripe of translation at this stage, using a transgene with the *nos* enhancer and promoter driving expression of scFv-NG. At the time, in the interpretation of our imaging data for zygotic *24xSunTag-hb* mRNAs, we had discussed whether the amount of scFv-NG protein was limiting, but disfavoured this interpretation based on the translation stripe persisting after we introduced extra copies of the scFv-NG transgene (Vinter et al., 2021a). However, since we detect uniform translation of zygotic *24xSunTag-hb* mRNAs in early nc14 embryos when we use the GAL4-UAS system to express higher levels of the scFv-sfGFP-NLS proteins here (Fig. S3A-C), we conclude that the scFv-NG proteins had become limiting in our previous experiments.

**Fig 3.**
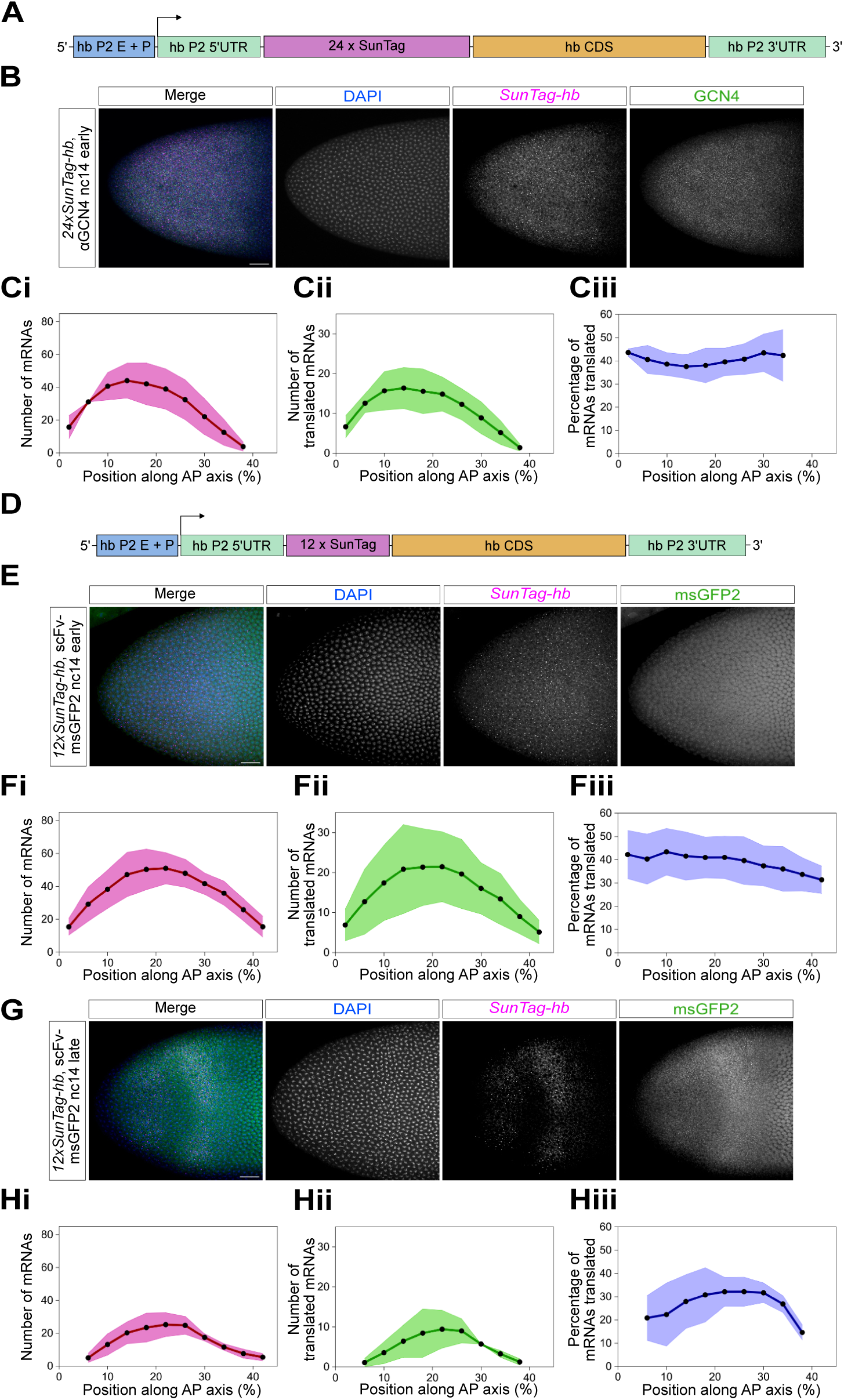
Zygotic *24xSunTag-hb* mRNA translation in nc14. (A) Schematic of the *hbP2>24xSunTag-hb* transgene. hb P2 E + P = hb P2 enhancer and promoter. (B) Image shows the zygotic *24xSunTag-hb* expression domain in an early nc14 embryo from a female carrying the *hbP2>24xSunTag-hb* transgene, stained with anti-GCN4 antibody (green), DAPI (blue) and *SunTag* smFISH probes (magenta). Single channel images are shown for clarity. Scale bar: 25 μm. (C) Quantitation of (i) total number of mRNAs, (ii) translated mRNAs and (iii) the percentage of mRNAs translated, per binned nuclear territory for 3 biological repeats, including the embryo shown in (B). Nuclear territories are shown as 20 μm bins along the AP axis. Data shown as mean value for each bin ± s.d. (D) As in (A) for the *hbP2>12xSunTag-hb* transgene. (E, G) Image of the zygotic *12xSunTag-hb* expression domain in early (E) and late (G) nc14 embryos, collected from *nos-GAL4-*VP16/*UASp-scFv-msGFP2-NLS* females crossed to *hbP2>12xSunTag-hb* males. Images show msGFP2 signals (green), DAPI staining (blue) and *12xSunTag-hb* mRNAs detected by smFISH probes (magenta). (F, H) Quantitation of (E) and (G) respectively, as described in (C).

To further investigate the translation profile at nc14, we used a GCN4 antibody, which recognises the SunTag epitopes, and SunTag smFISH probes to detect translation sites in fixed embryos carrying the *24xSunTag-hb* transgene. These data again show a constant proportion of *24xSunTag-hb* mRNAs translated across the expression domain in early nc14 embryos (Fig. 3B, C, Fig. S3G). We also find mostly uniform translation efficiency at late nc14, using the GCN4 antibody (Fig. S3H-J). At this stage the mRNA pattern is refining and a lower percentage of mRNAs are being translated (Fig. S3H-J). In contrast, detection of translation in embryos with *matα4-GAL4-VP16* driving expression of *UASp-scFv-sfGFP-NLS* reveals a stripe at the posterior of the expression domain (Fig. S3D-F). Analysis of embryos from females expressing *nos-GAL4-VP16* and *UASp-scFv-msGFP2-NLS* also revealed that scFv-msGFP2 proteins can be depleted in late nc14 embryos (Fig. S3K-M). Together, these data suggest that, even though scFv-sfGFP had been highly expressed using GAL4 amplification so that it is no longer limiting at early nc14, the protein has been depleted at late nc14.

We next generated a zygotic *12xSunTag-hb* transgene (Fig. 3D) to test whether the reduced number of SunTag repeats would be sufficient to allow scFv-FP detection of translation sites, while potentially avoiding depletion of scFv-FP proteins in late nc14 embryos. Embryos were collected from *nos-GAL4-VP16*/*UASp-scFv-msGFP2-NLS* females crossed to males carrying the *12xSunTag-hb* transgene. Translation sites are detectable in embryos, although the signals are weaker due to the reduced number of SunTag repeats (Fig. 3E, G). In early nc14 embryos, translation is uniform across the expression domain (Fig. 3F, Fig. S3N). At late nc14, the proportion of mRNAs translated is also mostly uniform (Fig. 3H, Fig. S3O), as observed for embryos with *24xSunTag-hb* mRNAs stained with anti-GCN4, indicating that reducing the copies of SunTag repeats to 12 avoids depletion of scFv-msGFP2 proteins. Together, these experiments highlight that scFv-FP proteins can be depleted for abundant, highly translated mRNAs with 24 or more SunTag copies, as also described recently by others (Bellec et al., 2024; Chen et al., 2024). Furthermore, using a 12xSunTag array with GAL4 driven expression of scFv-FP can be a useful strategy for ensuring that scFv-FP levels do not become limiting.

### Ribosome pausing *in vivo*

Ribosomes can pause during mRNA translation, with the *xbp1* mRNA a paradigm for studying ribosome pausing that is conserved from yeast to humans (Chyżyńska et al., 2021; Yan et al., 2016; Yanagitani et al., 2009; Yanagitani et al., 2011). Therefore, we used the *xbp1* mRNA ribosome pause site to test whether the zygotic *24xSunTag-hb* transgene could be used as a reporter to study ribosome pausing during development. Ribosome profiling and RNA-seq reads for *xbp1* are shown from published data for 0-2 hr *Drosophila* embryos (Dunn et al., 2013). The ribosome profiling data show an accumulation of reads at the site defined as the *xbp1* pause site in yeast and human tissue culture cells (Chyżyńska et al., 2021; Yanagitani et al., 2011), indicating that this pause site is functional in the early embryo (Fig. 4A). We inserted the *xbp1* pause site at the end of the *hb* coding sequence in the zygotic *24xSunTag-hb* transgene (Fig. 4B). This location was chosen so that a potential queue of ribosomes caused by pausing would not be restricted by lack of space on the mRNA (Goldman et al., 2021).

**Fig 4.**
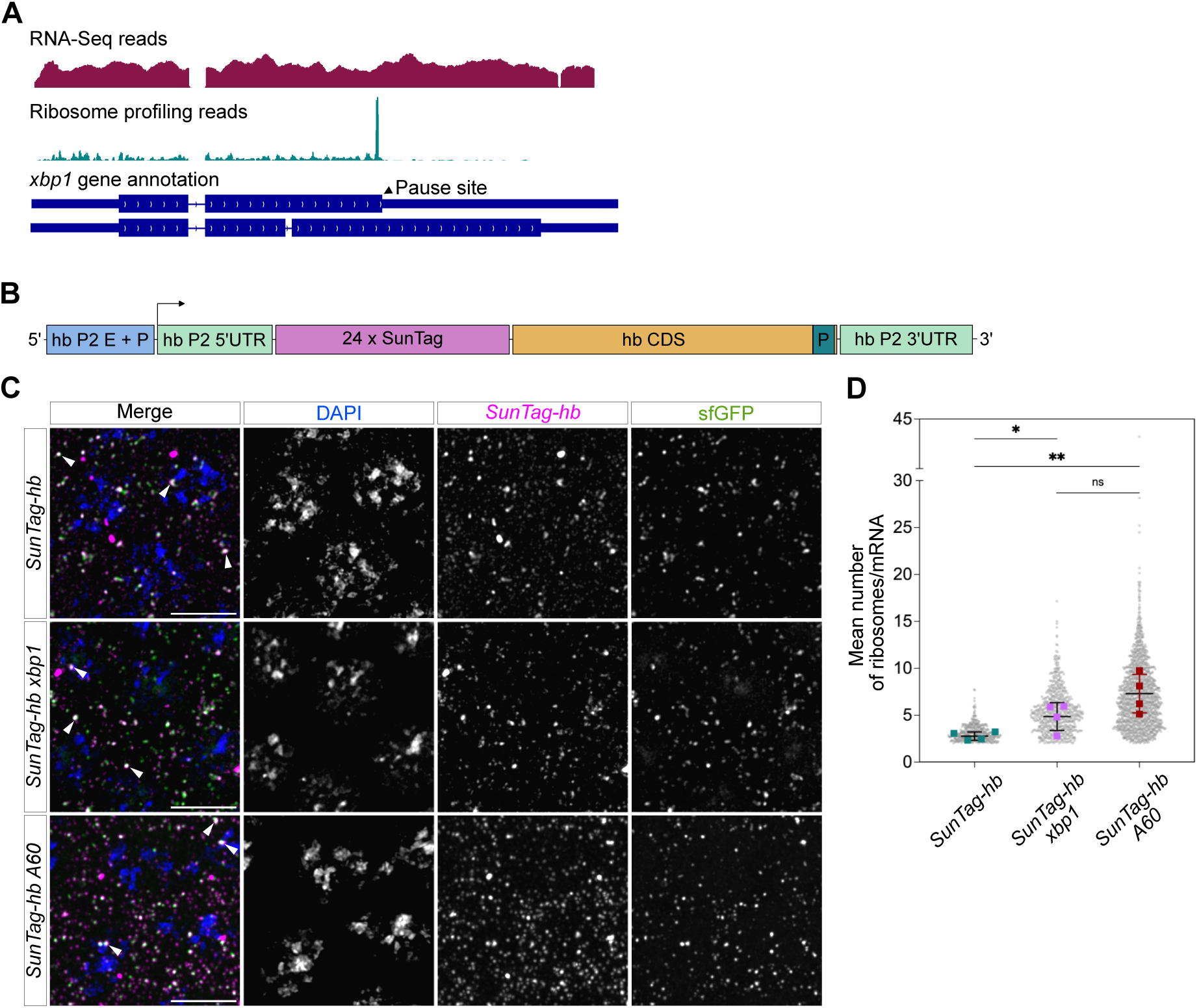
Quantitation of ribosome pausing in the embryo. (A) Genome browser view showing RNA-seq reads and ribosome profiling reads for *xbp1* in 0-2h *Drosophila* embryos. Data from (Dunn et al., 2013). (B) Schematic of the *hbP2>24xSunTag-hb* transgene with the pause site (P), either *xbp1* or A60, inserted upstream of the stop codon. hb P2 E + P = hb P2 enhancer and promoter. (C) Representative high magnification images of the anterior region of early nc14 embryos from *matα4-GAL4-VP16 UASp-scFv-sfGFP-NLS* females crossed to *hbP2>24xSunTag*-*hb*, *hbP2>24xSunTag-hb-xbp1* or *hbP2>24xSunTag-hb-A60* males. Images show sfGFP signals (green), *SunTag* smFISH staining (magenta) and DAPI (blue). White arrowheads show translation sites based on colocalised *SunTag-hb* and scFv-sfGFP signals. Scale bars: 5 μm. (D) Graph shows all the individual data points and mean numbers, from 4 biological replicates, of ribosomes present on zygotic *24xSunTag*-*hb*, *24xSunTag-hb-xbp1* and *24xSunTag-hb-A60* males, unpaired two-tailed *t-*test, ns not significant, \**P*<0.05, \*\**P*<0.01. Data are mean ± s.d.

Embryos from *matα4-GAL4-VP16 UASp-scFv-sfGFP-NLS* females crossed to homozygous *24xSunTag-hb-xbp1* males were collected and mRNAs and translation sites were detected using smFISH and sfGFP fluorescent signals, respectively. An image from the anterior region of a representative embryo at early nc14, a stage when scFv-sfGFP is not limiting, is shown in Fig. 4C. Ribosome number/mRNA was quantitated, as described above, for *24xSunTag-hb-xbp1* mRNAs relative to *24xSunTag-hb* mRNAs. Introducing the *xbp1* pause site results in a significant increase (∼2-fold) in the mean number of ribosomes on *24xSunTag-hb* mRNAs in the anterior region of early nc14 embryos (Fig. 4D). We also tested the effect of inserting a 60-nucleotide poly(A) sequence at the same position in the *SunTag-hb* transgene. Poly(A) has been shown to induce ribosome pausing in mammalian cells (Chandrasekaran et al., 2019; Koutmou et al., 2015; Tesina et al., 2020), including in an equivalent SunTag pausing reporter in human tissue culture cells (Goldman et al., 2021). In the anterior of early nc14 embryos, A60 also induces a significant increase (∼3-fold) in ribosome number, consistent with ribosome pausing (Fig. 4D).

We also analysed the anterior regions of nc12 and nc13 embryos in the same way to determine whether the extent of ribosome pausing changed over developmental time. These data show that ribosome pausing is not detected for *24xSunTag-hb-xbp1* or *24xSunTag-hb-A60* mRNAs at nc12 (Fig. S4A, B). However, in nc13 embryos, similar increases in ribosome number to that observed at nc14 are detected in the presence of the *xbp1* or *A60* pause sequences (Fig. S4C, D). Together, these data show that the *24xSunTag-hb* transgene can be used as a reporter to evaluate and study potential ribosome pausing sequences during development. In addition, these findings raise the possibility that pausing may be developmentally regulated.

### Visualisation and quantitation of *sog* mRNA translation in fixed embryos

Next, we addressed whether the SunTag system can be used to visualise translation of mRNAs encoding secreted proteins. To this end, we studied the *short gastrulation* (*sog*) mRNA, encoding a secreted protein that binds to BMP signalling molecules extracellularly to form the BMP gradient required for dorsal-ventral axis patterning (Montanari et al., 2022). We generated a *LS>SunTag-sog* transgene, with transcription of *SunTag-sog* under the control of the endogenous *sog* promoter and lateral stripe (LS) enhancer (Markstein et al., 2002). As insertion of SunTag sequences N-terminal to the signal peptide would likely interfere with protein secretion (Pool, 2022), 24 copies of the SunTag peptide were added downstream of the *sog* signal peptide (Francois et al., 1994) (Fig. 5A). In this way the SunTag array will be positioned at the Sog N-terminus following cleavage of the signal peptide during translocation. After the signal peptide is translated, the ribosome is arrested until the signal peptide docks at an ER translocon (Pool, 2022).

**Fig 5.**
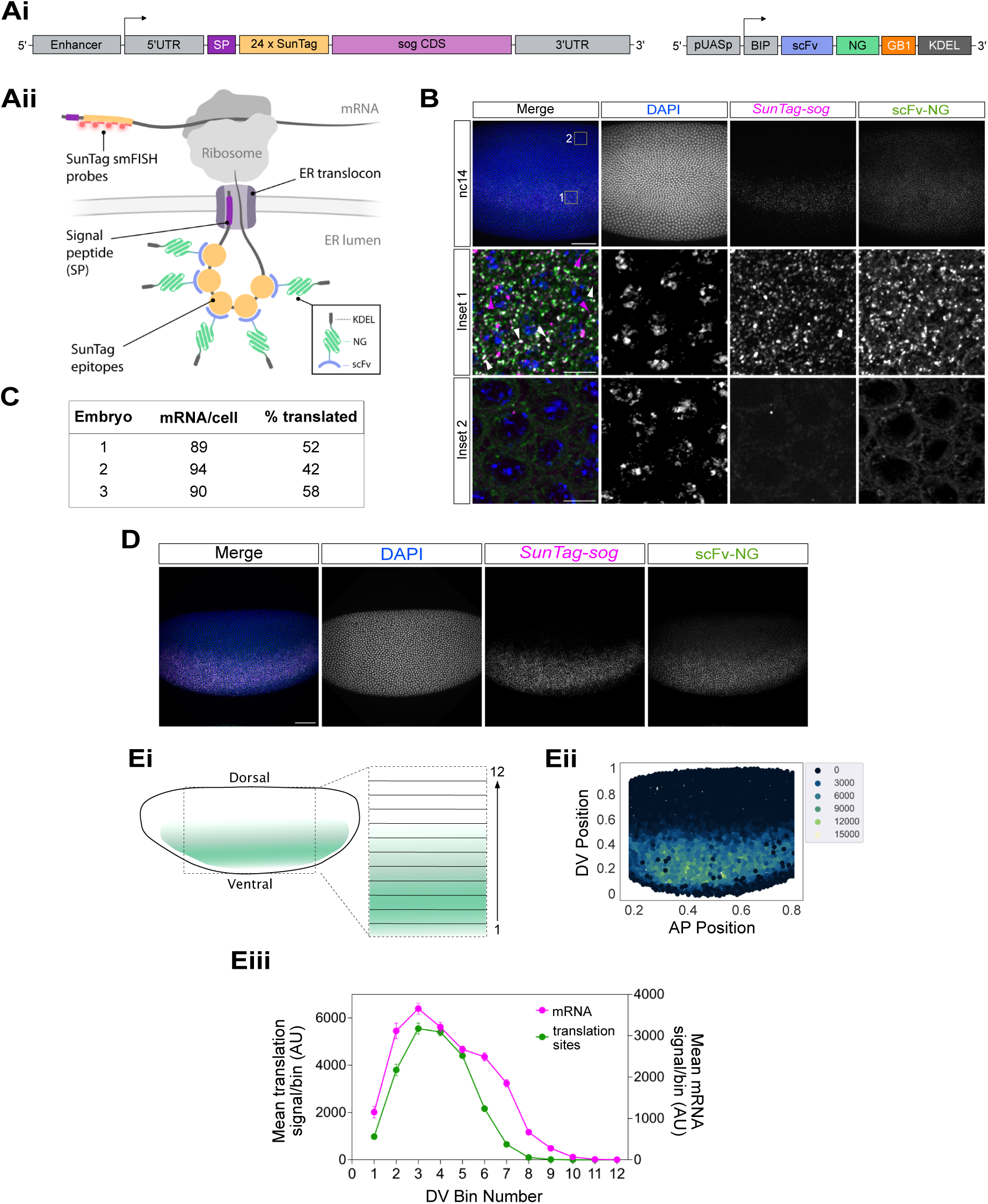
Visualising translation of the secreted Sog protein with the SunTag system. (A) (i) Schematics of the LS>*SunTag-sog* and *UASp-BiP-scFv-NG-GB1-KDEL* transgenes used to visualise Sog translation. *sog* transcription is under the control of the lateral stripe (LS) enhancer and promoter, SP indicates signal peptide. (ii) Schematic showing detection of *SunTag*-*sog* mRNA translation. The mRNA is detected by SunTag smFISH probes. After the signal peptide docks at the ER translocon, the translated SunTag epitopes are recognised by scFv-NG-KDEL fusion proteins resident in the ER. (B) Central region of a fixed nc14 embryo (lateral view) from females carrying *nos-GAL4-VP16* and *UASp*-*BIP-scFv-NG-KDEL* insertions crossed to *LS>SunTag-sog* males, showing NG signal (green) and stained with DAPI (blue) and *SunTag* smFISH probes (magenta). The embryo areas used for the high magnification images are shown. White arrowheads indicate translation sites, detected as clusters of colocalised scFv-NG-KDEL and mRNA signals within the expression domain. Magenta arrowheads indicate single mRNAs, whereas the translated *SunTag-sog* mRNAs are clusters of multiple mRNAs. Scale bar: 50 μm for lateral view, 5 μm for insets. (C) Table showing the quantitation of mRNA number and proportion translated for 3 biological repeat embryos, including the one shown in (B). (D) Images show a lateral view of a fixed nc14 embryo as described in (B). Scale bar: 50 μm. (E) (i) Cartoon of a lateral view of the *sog* expression domain, with 12 bins across the DV axis. The box indicates the region of analysis. (ii) Heat map showing the total NG signal per cell. (iii) Quantitation of the mean total NG and mRNA signals per binned cell. Cells are grouped into the 12 bins across the DV axis and data is shown as the mean value of each bin ± S.E.M.

Embryos carrying only the *LS>SunTag-sog* transgene show apical localisation of the *SunTag-sog* mRNAs (Fig. S5A), as described for *sog* mRNAs previously (Reeves et al., 2012), indicating that insertion of the SunTag sequences does not affect *sog* mRNA localisation. Using antibody staining with the GCN4 antibody and smFISH with SunTag probes, we detected *SunTag-sog* mRNA translation (Fig. S5B). However, using the experimental set up of collecting embryos from *nos>scFv-NG* females crossed to *SunTag-sog* transgenic males, no translation sites were detected in embryos. This is consistent with translation being arrested in the cytoplasm after the signal peptide, so that the SunTag array is translated at the ER and inserted into the lumen where it is inaccessible to cytoplasmic scFv-NG proteins.

To promote ER targeting of the scFv-NG proteins, we next generated a fly stock with a transgene containing scFv-NG downstream of a signal sequence so that it would be secreted. However, translation sites were still undetectable in embryos from females maternally expressing this secreted scFv-NG protein and carrying the *SunTag-sog* transgene. We reasoned that this was because the scFv-NG protein was efficiently secreted and therefore levels in the ER were too low to detect the SunTag peptides being translated. Therefore, we included a KDEL ER retention signal to generate a BiP-scFv-NG-GB1-KDEL transgene (hereafter called BiP-scFv-NG-KDEL) (Fig. 5Ai). The GB1 solubility tag that was included (Tanenbaum et al., 2014) is present in all other ScFv-FP fusions used in this study. The BiP signal sequence allows efficient ER targeting in *Drosophila* cells (Iwaki and Castellino, 2008), whereas the presence of the KDEL sequence ensures ER retention (Newstead and Barr, 2020) so that the scFv-NG protein can recognise the newly translated SunTag peptides (Fig. 5Aii).

Embryos were collected from females carrying *nos-GAL4-VP16* and *UASp-BiP-scFv-NG-KDEL* insertions crossed to either control males or those homozygous for the *LS>SunTag-sog* transgene. mRNAs were detected by smFISH with SunTag probes and translation of the SunTag peptides was detected by NG fluorescence (Fig. 5B). In nc14 control embryos lacking the *LS>SunTag-sog* transgene, weak NG signals were visible in cells in the neuroectoderm and dorsal ectoderm (Fig. S5C), in a pattern consistent with ER localisation (Kilwein and Welte, 2021). Similar staining was detected in *LS>SunTag-sog* nc14 embryos outside the *sog* expression domain (Fig. 5B, inset 2). In the neuroectoderm where *sog* is expressed (Francois et al., 1994), much stronger NG signals are detected that are colocalised with the mRNA signals (Fig. 5B, inset 1). As documented previously, *sog* mRNAs are mostly present in clusters rather than as individual mRNAs (Frampton et al., 2022), likely due to their association with the ER for translation.

We used these high resolution images to estimate the number of *SunTag-sog* mRNAs per cell and proportion translated. As the *SunTag-sog* mRNAs are mostly in clumps, it is not possible to simply count individual mRNAs. Instead, we quantitated the total amount of *SunTag-sog* mRNA smFISH signals and divided by the mean signal from the small number of individual *SunTag-sog* mRNAs detected (Fig. 5B). This value was divided by the number of cells in the image to give an approximation of the number of *SunTag-sog* mRNAs per cell (see Methods). The data for 3 biological replicates suggest that there are ∼90 *SunTag-sog* mRNAs/cell with ∼50% of these translated (Fig. 5C).

As the above analysis focused on only a small region of the *SunTag-sog* expression domain, we used whole embryo images to quantitate *SunTag-sog* mRNA translation sites across the expression domain (Fig. 5D). To this end, the *SunTag-sog* expression domain was divided into 12 bins across the DV axis in the centre of the embryo (Fig. 5Ei). Translation sites were called based on NG signals being colocalised with clustered mRNA signals, then assigned to the closest nucleus (see Methods). Based on this, the total translation signal per (virtual) cell was quantitated and is shown as a heatmap for a representative nc14 embryo (Fig. 5Eii). In addition, the mean translation signal data for all the cells in each of the DV bins is shown in Fig. 5Eiii. In these whole embryo images, it is not possible to discern the small number of single *SunTag-sog* mRNAs. Instead, the mean total mRNA signal was quantitated, which shows that the translation profile mirrors that of the *SunTag-sog* mRNAs in each embryo (Fig. 5Eiii). The same trend is observed for data from 2 other biological repeat embryos (Fig. S5D-G). Together, these data from fixed embryos suggest that there is relatively uniform translation (∼50%) of *SunTag-sog* mRNAs in nc14 embryos.

### Live imaging of *sog* mRNA translation

In addition to imaging *SunTag-sog* mRNA translation in fixed embryos, we tested scFv-NG-KDEL detection in live imaging. *nos-GAL4-VP16*/*UASp-BiP-scFv-NG-KDEL* females were crossed to *LS>SunTag-sog* or control males and embryos were collected and imaged live during early development. In these embryos, there is weak scFv-NG-KDEL signal localised to the ER, so that the nuclei can be discerned based on their absence of fluorescent signal (Fig. 6A). This allowed us to age the embryos across the early cleavage cycles. Stills from a live imaging movie of a control nc14 embryo show background fluorescence signals in the neuroectoderm (Fig. S6A). Quantitation of the background fluorescence in the neuroectoderm of 3 biological repeat control embryos reveals that it is uniform across developmental time (Fig. S6B).

**Fig 6.**
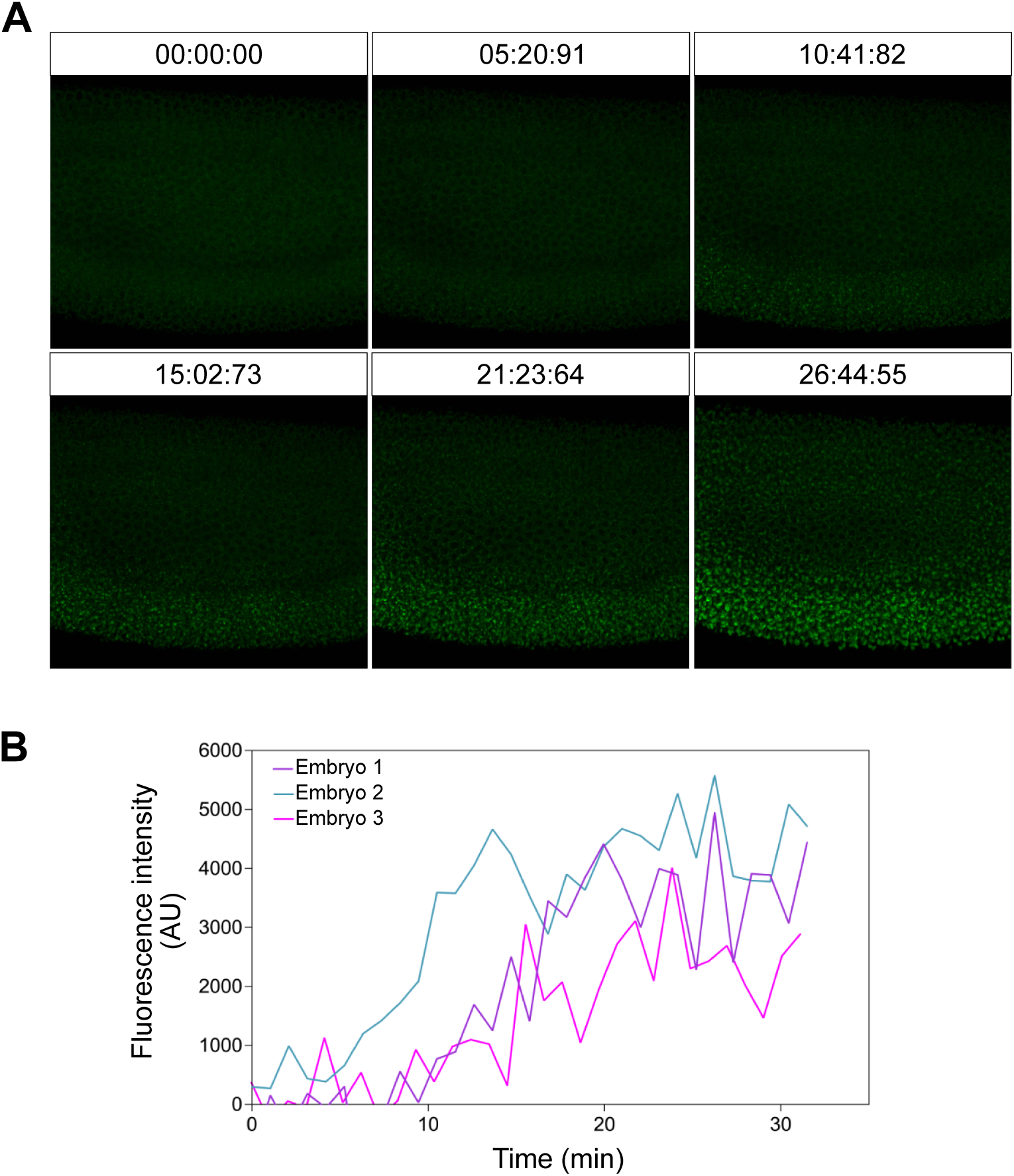
Live imaging of *SunTag*-*sog* mRNA translation. (A) Stills from a movie of a *SunTag-sog* embryo developing through nc14, with translation detected by scFv-NG-KDEL proteins. The embryo was collected from females carrying *nos-GAL4-VP16* and *UASp*-*BIP-scFv-NG-KDEL* crossed to *SunTag-sog* males. Time intervals are min:s:ms. See Movie 1. (B) Quantitation of NG signals in the neuroectoderm over a ∼30 min period in nc14. Data are from 3 biological replicates.

In contrast, stills from a movie capturing 30 min of nc14 show that NG translation site signals accumulate in the neuroectoderm where *SunTag-sog* is expressed (Fig. 6A). Quantitation of the total fluorescence intensity in the neuroectoderm of 3 biological repeat embryos reveals that translation sites increase throughout nc14 (Fig. 6B). Together, the data for *SunTag-sog* show that the scFv-NG-KDEL system can be used to visualise translation of *SunTag-sog* mRNAs in both fixed and live embryos, facilitating the study of translation of other mRNAs encoding secreted proteins.

## Discussion

A major aspiration in the field of protein synthesis has been to study the process of translation and its regulation at single mRNA resolution. Recent advances in fluorescent microscopy techniques have enabled such studies to be conducted. For instance, the SunTag imaging method enables the detection of nascent proteins on single mRNAs such that ribosome numbers and rates of elongation can be estimated (Pichon et al., 2016; Wang et al., 2016; Wu et al., 2016; Yan et al., 2016). In this study, we explore the scope of this SunTag method in precisely staged *Drosophila* embryos to provide a developmental context to the study of translation. More specifically, we extend the use of the SunTag method in the *Drosophila* embryo to study the translation of maternal and zygotic transcripts, investigate ribosome pausing and examine the production of secreted proteins at the ER membrane.

We show uniform translation of zygotic *SunTag-hb* mRNAs in nc14 embryos, but depletion of scFv-FP proteins is a problem, even for *24xSunTag-hb* mRNAs in late nc14 embryos when GAL4 amplification is used to express high scFv-FP levels. Limiting scFv-FP proteins has also been described in other recent SunTag studies in *Drosophila* (Bellec et al., 2024; Chen et al., 2024). Our data show that including 12x SunTag repeats overcomes scFv-FP depletion.

Although there is a trade-off of slightly lower signal to noise, our estimates of the proportion of translated *12xSunTag-hb* mRNAs are similar to those from GCN4 staining or the *24xSunTag-hb* reporter when scFv-FP is not limiting. Likewise, in mammalian cells, comparable translation elongation rates and ribosome densities were obtained using a reporter with either 5x or 24x SunTag repeats (Yan et al., 2016). Therefore, in addition to control GCN4 antibody staining, testing a 12x SunTag insertion will serve as a useful control to avoid limiting scFv-FP proteins. The smaller 12x cassette also has the advantage that it may be less likely to affect mRNA stability, translation and/or protein function.

The maternal *hb* mRNA has served as a paradigm for the study of translation repression during development, with this repression essential for correct abdominal patterning (Hülskamp et al., 1989; Irish et al., 1989; Struhl, 1989). By visualising and quantitating translation sites, we were able to directly measure the strength of translational repression in the embryo. We find that ∼3-5% of the maternal *SunTag-hb* mRNAs are weakly translated in the posterior of the early embryo compared to ∼50-60% in the anterior. As we saw a similar very low proportion of *SunTag-hb* mRNAs translated in the posterior of embryos with 1 or 2 copies of the maternal *SunTag-hb* transgene, we consider it unlikely that this is due to titration of a repressor, but a CRISPR insertion in the endogenous *hb* gene is required to fully test this. Other examples of incomplete translation repression have been uncovered by SunTag imaging. In the early *Drosophila* embryo, *nanos* mRNAs are translated in germ granules but repressed in the soma. *SunTag-nanos* imaging has revealed that ∼30-50% of *SunTag-nanos* mRNAs are translated in the germ plasm, compared to less than 2% in the soma (Chen et al., 2024). In mammalian cells, SunTag imaging of a reporter with the long 5’UTR of the Early mitotic inhibitor 1 gene revealed that although the mRNAs are mostly strongly translationally repressed, ∼2% escape translation repression and are robustly translated (Yan et al., 2016).

The Pumilio (Pum), Nos and Brain Tumor (Brat) RNA binding proteins bind to two Nos-response elements in the 3’ UTR of the maternal *hb* mRNA to repress its translation and promote its decay in the posterior of the embryo (Arvola et al., 2017). Based on evidence that Brat and Pum-Nos interact with the *hb* mRNA independently, it has been suggested that separate repression mechanisms may exist (Macošek et al., 2021). Consistent with this, Brat recruits 4EHP, which binds to the cap and inhibits translation, although some translation repression of the *hb* mRNA was still observed in the posterior of 4EHP hypomorphic mutant embryos (Cho et al., 2006). Future studies can exploit the maternal *SunTag-hb* transgene described here and extensive collection of well-characterised Pum, Nos and Brat mutants (Arvola et al., 2017) to quantitate how loss of specific interactions affect the efficiency and strength of translation repression. Moreover, as Nos is targeted to thousands of maternal mRNAs during oogenesis, leading to their inefficient translation (Marhabaie et al., 2024), the SunTag system can be used to uncover the mechanism of this regulation of the maternal transcriptome.

mRNA UTRs can influence the translation initiation rate (Leppek et al., 2018; Mayr, 2017). However, we find that, despite having different UTRs (Margolis et al., 1995; Schröder et al., 1988), maternal and zygotic *SunTag-hb* mRNAs have similar numbers of ribosomes at nc12 and nc13. Ribosome number on zygotic *SunTag-hb* mRNAs is similar at nc12 and nc13, then declines in early nc14 embryos. In the *Drosophila* embryo, zygotic genome activation occurs in two waves: a minor wave from nc8-13 in which transcription of ∼100 genes is activated, followed by a major wave at nc14 that involves the activation of thousands of genes (Hamm and Harrison, 2018). We speculate that the large increase in numbers of zygotic transcripts at nc14 results in a greater competition for the ribosome pool or limiting translation factors, so that ribosome density on zygotic *SunTag-hb* mRNAs decreases.

Little is known about ribosome pausing during development. Here we show that the *xbp1* pause site and an A60 sequence both lead to an increased number of ribosomes on the *SunTag-hb* mRNAs in nc13 and nc14 embryos, consistent with ribosome pausing. However, no pausing was detected with the pause site reporters in nc12 embryos, suggesting that pausing can be developmentally regulated. Analysis of *xbp1* translational pausing in mammalian cells identified a 26–amino acid region at the C-terminus of XBP1u. Alanine scanning mutagenesis showed that 14 amino acids in this region of XBP1u are important for pausing, whereas an S255A mutation strengthened the pause (Yanagitani et al., 2011). This enhanced pause site induced ribosome pausing in a SunTag reporter in human cells, with heterogeneity in the pause duration (Yan et al., 2016). The fly *xbp1* pause sequence is quite divergent in comparison to the highly conserved sequences of the human, zebrafish and mouse pause sites. For example, the codons in the P- and A-site of the paused ribosome are different in flies, and the fly sequence also contains an A at the equivalent position to human S255 that was found to increase pausing (Chyżyńska et al., 2021). In future, the *SunTag-hb-xbp1* reporter could be used to dissect the sequence requirements for the fly *xbp1* pause site.

Inappropriate polyadenylation at sites within the coding region of mRNAs is one of the more common defects found in eukaryotic mRNAs (Ozsolak et al., 2010). Such mRNAs with their defective poly(A) sequences cause translating ribosomes to pause to initiate pathways of mRNA degradation, nascent protein destruction and ribosome recycling (Höpfler and Hegde, 2023). The mechanics of poly(A)-dependent ribosome pausing are thought to rely upon a combination of the impact of the encoded poly-lysine on the ribosomal peptidyl transfer reaction and the effects of a stable helical structure formed by poly(A) in the mRNA decoding centre (Chandrasekaran et al., 2019). The propensity for inappropriate polyadenylation and therefore the role of this pathway in developmental contexts is largely unknown. In this study, we show that a *SunTag-hb-A60* reporter can be used to study poly(A) dependent ribosome pausing in the whole *Drosophila* embryo. This opens up the possibility of using this system in combination with genetic strategies to study the mechanics of removal of inappropriately polyadenylated mRNAs in a whole living organism.

In a final application of the SunTag system, we use a *LS>SunTag-sog* transgene to study translation of a secreted protein. For this, targeting the scFv-NG protein for secretion was insufficient to allow detection of translation sites, with a KDEL ER retention signal also being required. While this allows *SunTag-sog* mRNA translation sites to be visualised, it appears that the translated SunTag-Sog proteins (with bound scFv-NG-KDEL) are also ER retained, although additional experiments are required to fully address this. Therefore, if CRISPR editing is used to insert SunTag sequences into the gene of interest encoding a secreted protein, studying translation in embryos heterozygous for the SunTag insertion would avoid lethality in the presence of scFv-NG-KDEL proteins. In addition, it is difficult to estimate ribosome number as the clustering of mRNAs at the ER and protein retention mean that it is challenging to detect single mRNAs and nascent proteins. The auxin-inducible degron was added to the C-terminus of a SunTag reporter in mammalian cells, to reduce the fluorescent background by allowing fully translated proteins to be degraded (Wu et al., 2016). A similar optogenetic approach with the blue light-inducible degron (Irizarry et al., 2020) could be used here to reduce background from ER-retained fully translated SunTag-Sog proteins, or the scFv-NG-KDEL protein could be expressed in a pulsed manner, e.g. using a heatshock promoter (Lis and Wu, 1993).

The *sog* gene is very long (∼30 kb), with a truncated form of the mRNA detected in nc13 due to use of an alternative poly(A) signal. It has been suggested that this shorter form overcomes the time constraints associated with transcribing a long mRNA in the short early nuclear cleavage cycles (Sandler et al., 2018), but the translation rate will also be important. Translation elongation rates in *Drosophila* measured using the SunTag system and FRAP or FCS approaches range from 4-35 amino acids/second (Chen et al., 2024; Dufourt et al., 2021). If individual translation sites can be visualised, future studies can measure the translation rates of *SunTag-sog* mRNAs across the early cleavage cycles to determine the extent to which this is tuned to developmental timing.

In summary, the SunTag quantitative imaging method holds great promise for the study of mRNA translation in development. Its impact is expected to be similar to that of quantitative imaging of transcription in *Drosophila,* which has uncovered new concepts relating to gene regulation (Fukaya et al., 2016). Overall, the results described here show the versatility of the SunTag approach and will facilitate the study of new facets of translation regulation during development.

## Materials and Methods

### Cloning

#### phbP2>24xSunTag-hb-xbp1, phbP2>24xSunTag-hb-A60, phbP2>12xSunTag-hb and pMatE>24xSunTag-hb

phbP2>24xSunTag-hb has been described (Vinter et al., 2021a). phbP2>24xSunTag-hb-xbp1 was made using phbP2>24xSunTag-hb as a template, with a hbP2>24xSunTag-hb fragment and an xbp1 fragment (2R:21144772-21144862) inserted using 2-way In-Fusion into StuI/BamHI digested phbP2>24xSunTag-hb. phbP2>24xSunTag-hb-p60A was generated by inserting hbP2>24xSuntag-hb (from phbP2>24xSunTag-hb) and p60A (encoded by an oligo) into hbP2>24xSunTag-hb digested with StuI/BamHI. For pMatE>SunTag-hb, the *hb* CDS and *hb* 3’UTR were subcloned from phbP2>24xSunTag-hb into pAc5.1A. The first 18 bases of *hb* exon 2, the start codon and a NotI site were introduced using annealed oligos. The resulting hb exon2-NotI-hbCDS-3’UTR was then amplified and the maternal *hb* enhancer and *hb* P1 exon 1 (amplified form genomic DNA) were inserted into the HindIII/NdeI site of pUASp-attB (DGRC_1358, RRID:DGRC_1358) by multiple insert In-Fusion (Takara Biosciences) cloning. The SunTag was then inserted into the NotI site. To generate hbP2>12xSuntag-hb, hbP2>12xSuntag and hb CDS were amplified from phbP2>24xSuntag-hb and inserted into StuI/BamHI digested phbP2>24xSuntag-hb using In-Fusion cloning. See Table S1 for primers.

#### pUASp-BiP-scFv-mNeonGreen-GB1-KDEL-tubulin 3’UTR

To generate pCasper-nos>BiPscFv-mNeonGreen-GB1-KDEL-tubulin 3’UTR, mNeonGreen-GB1-KDEL was first amplified from pCasper-nos>scFv-mNeonGreen-GB1-NLS (Vinter et al., 2021a) with a KDEL sequence in the primer and inserted back into the BamHI site of pCasper-nos>scFv-mNeonGreen-GB1-NLS-tubulin 3’UTR (Vinter et al., 2021a) using In-Fusion. Then, BiP-scFv-mNeonGreen-GB1-KDEL-tubulin 3’UTR was amplified, with a primer containing the BiP secretion sequence, and cloned into the NotI/XbaI sites of pUASp-attB using In-Fusion.

#### RIV-white-LS>SunTag-sog

The lateral stripe (LS) enhancer (Markstein et al., 2002), promoter region, 5’UTR and CDS were subcloned from pPelican-ls-sog (Peluso et al., 2011) with the sog 3’UTR (amplified from genomic DNA) into RIV-white (DGRC 1330, RRID:DGRC_1330). SunTag (from pcDNA4TO-24xGCN4_v4_sfGFP (Addgene #61058, RRID:Addgene_61058) (Tanenbaum et al., 2014) was inserted downstream of the transmembrane domain in the sog CDS flanked by two linkers consisting of 4xGly 1xSer 4xGly. All plasmids are available from the Ashe lab on request.

### Fly Stocks

Fly lines were raised with standard fly food mix (glucose 78g/l, maize flour 72g/l, yeast 50g/l, agar 8g/l, Nipagin 27ml/l and propionic acid 3ml/L) and maintained at 18°C. All experimental crosses were performed at 25°C. Fly stocks used are listed in Table 1. Transgenic lines were made by the University of Manchester microinjection service. The line *y^1^w*;;UASp-scFv-sfGFP-NLS/TM3Sb* was recombined with *w*;;P{matα4-GAL-VP16}V37* to generate w*;;*P{matα4-GAL-VP16}V37 UASp-scFv-sfGFP-NLS/TM6B*, although we note that a proportion of embryos from these females arrest development. For imaging zygotic *24xSunTag-hb* or *12xSunTag-hb* mRNAs, embryos were collected from *w*;;P{matα4-GAL-VP16}V37 UASp-scFv-sfGFP-NLS/TM6b* or *nos-GAL4-VP16/UASp-scFv-msGFP2-NLS* females crossed to *hbP2>24xSunTag-hb* or *hbP2>12xSunTag-hb* males. For the GCN4 staining, embryos were collected from *y^1^w^67c23^* females crossed to *hbP2>24xSunTag-hb* males.

**Table 1.**
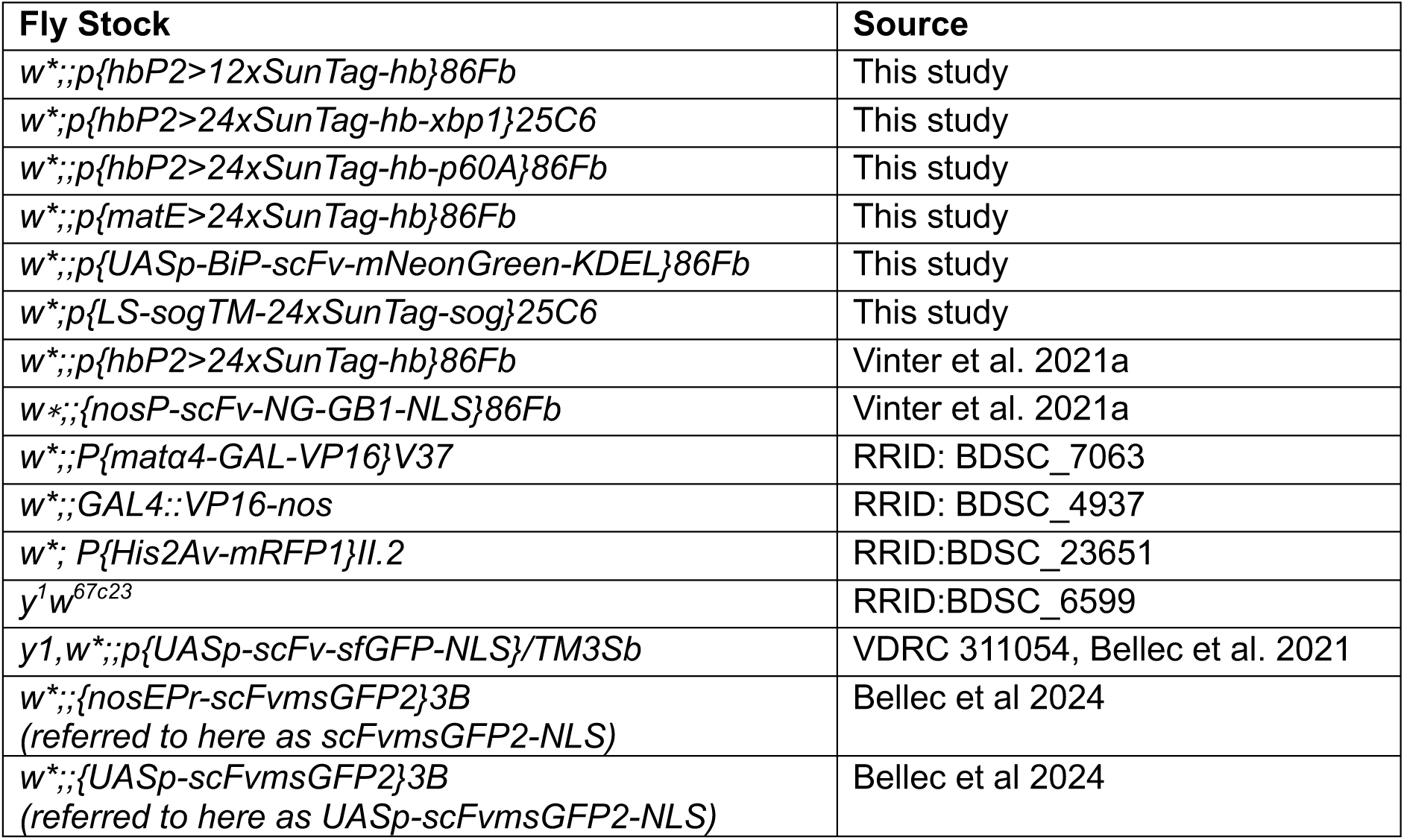
Fly stocks used in this study.

To study maternal *24xSunTag-hb* mRNAs, *w*;;P{matα4-GAL-VP16}V37 UASp-scFv-sfGFP-NLS/matE>24xSunTag-hb* females were crossed to *y^1^w^67c23^*males for embryo collections. For the GCN4 staining with the maternal transgene, embryos were collected from *matE>24xSunTag-hb*/+ (for 1 copy) or *matE>24xSunTag-hb* (for 2 copies) females. For *LS>SunTag-sog* imaging, *w^∗^;; nos-GAL4-VP16*/*UASp-BiP-scFv-NG-KDEL* females were crossed to *w*;LS>SunTag-sog* or *y^1^w^67c23^* (for control embryos) males. Fly stocks are available from the Ashe lab on request. For the GCN4 staining of *LS>SunTag-sog* embryos, homozygous embryos were used.

### smFISH and immunofluorescence

*Drosophila* embryos (2-4h) were fixed as previously described (Vinter et al., 2021a) and stored in methanol at −20°C. Fixed embryos were transferred into glass scintillation vials and washed 5 min in 50% methanol/50% phosphate-buffered saline with 0.1% Tween-20 (Merck P1379) (PBT), followed by four 10 min washes in PBT, a 10 min wash in 50% PBT/50% wash buffer (10% formamide in 2X SSC), and two 5 min washes in 100% wash buffer. Embryos were then incubated twice for 30 min in smFISH hybridization buffer (2.5mM dextran sulphate, 10% formamide in 2X SSC) at 37°C. Quasar 570 *SunTag* smFISH probes (Biosearch Technologies) (Vinter et al., 2021a) were diluted in hybridisation buffer to a final concentration of 100 nM and embryos were incubated with probe solution for 14 h at 37°C. Embryos were washed for 30 min at 37°C with pre-warmed wash buffer before three additional 15 min washes at 37°C and a 15 min wash in the dark at room temperature. Embryos were rinsed twice with PBT, followed by four 15 min washes in PBT in the dark. The third wash included DAPI (1:500). When the GCN4 antibody was used, the embryos were blocked in 1x Western Blocking Reagent (WB) (Merck 11921673001) after the PBT rinses and then incubated overnight at 4°C with anti-GCN4 (C11L34) primary antibody (Novus Biologicals NBP2-81273, RRID:AB_3413163), 1:250 in 1xWB. Embryos were rinsed in PBT then four 15 min PBT washes were carried out before blocking for 30 min in 1xWB. Embryos were then incubated with secondary antibody (Donkey anti-mouse Alexa Fluor 488 Thermo Fisher Scientific Cat# A-21202, RRID:AB_141607), 1:250 in WB for 2 h at room temperature. Embryos were rinsed twice with PBT and then washed four times for 15 minutes with 1:500 DAPI in the third wash. Embryos were mounted in ProLong^TM^ Diamond Antifade Mountant (ThermoFisher Scientific P36961).

### Fixed embryo imaging

Images to analyse *SunTag-hb* mRNAs and translation sites were acquired on a Leica TCS SP8 AOBS inverted confocal microscope using a 40x/1.30 HC PL Apo CS2 oil objective with a 1.3x confocal zoom. The confocal settings were as follows: pinhole 1 airy unit, scan speed 600 Hz unidirectional, 3x line averaging and 2048×2048 pixel format. Images were illuminated with a white light laser at 70% and collected sequentially using either Photon Multiplying Tube Detectors or Hybrid Detectors. For scFv-sfGFP embryos the following detection mirror settings were used: Photon Multiplying Tube Detector for DAPI (5% 405 nm excitation, collection: 415-470 nm); Hybrid SMD Detectors for sfGFP (15% 488 nm excitation, collection: 498-540 nm, 1-6 ns gating) and Quasar 570 (20% 548 nm excitation, collection: 558-640 nm, 1-6 ns gating). For Alexa Fluor 488 antibody-stained embryos the following detection mirror settings were used: Photon Multiplying Tube Detector for DAPI (10% 405 nm excitation, collection: 415-470 nm); Hybrid Detectors for Alexa 488 (6.5% 488 nm excitation, collection: 498-540 nm, 1-6 ns gating) and Quasar 570 (30% 548 nm excitation, collection: 558-640 nm, 1-6 ns gating). For all samples 3D optical z-stacks were acquired with 300 nm spacing.

High magnification images for quantification of *SunTag-hb* mRNA ribosome numbers and quantitation of maternal *SunTag-hb* mRNA numbers and percent translated were acquired using a 100x/1.40 HC PL Apo CS2 oil objective with a 4x confocal zoom. The confocal settings were as follows, pinhole 0.65 airy unit, scan speed 600 Hz bidirectional, 6x line averaging and 2048×2048 pixel format. The following detection mirror settings were used: Photon Multiplying Tube Detector for DAPI (10% 405 nm excitation, collection: 415-470 nm); Hybrid Detectors for sfGFP (15% 488 nm excitation, collection: 498-540 nm, 1-6 ns gating) and Quasar 570 (30% 548 nm excitation, collection: 558-640 nm, 1-6 ns gating). 3D optical z-stacks were acquired with 200 nm spacing.

Images to analyse *SunTag-sog* mRNA translation sites were acquired on a Leica TCS SP8 AOBS inverted confocal microscope using a 40x/1.30 HC PL Apo CS2 oil objective with a 0.75x confocal zoom. The confocal settings were pinhole 1 airy unit, scan speed 600 Hz unidirectional, 3x line averaging and 2048×2048 pixel format. High magnification images of *SunTag-sog* mRNA translation sites were acquired using a 100x/1.40 HC PL Apo CS2 oil objective with a 6x confocal zoom and the same confocal settings. Images were illuminated with a white light laser at 70% and collected sequentially using either Photon Multiplying Tube Detectors or Hybrid SMD Detectors. The following detection mirror settings were used: Photon Multiplying Tube Detector for DAPI (10% 405 nm excitation, collection: 415-470 nm); Hybrid SMD Detectors for NG (2% 506 nm excitation, collection: 516-548 nm, 0.5-6 ns gating) and Quasar 570 (10% 548 nm excitation, collection: 558-640 nm, 0.5-6 ns gating). 3D optical z-stacks were acquired with 300 nm spacing for whole embryo images and 200 nm spacing for high magnification.

Raw images were deconvolved with Huygens Professional software (SVI). All images presented are max intensity projections, except for the except for the *SunTag-sog* images in Figures 5 and S5, which are projections of 10 slices. Note that in the max projection images, translation sites present above or below the nucleus can appear nuclear. All embryos are oriented with anterior to the left.

### Live imaging

Embryos were dechorionated in 50% bleach for 2 min and mounted onto a Lumox imaging dish (Sarstedt, 94.6077.305) (Hoppe and Ashe, 2021; Vinter et al., 2021b). To visualise *SunTag-sog* mRNA translation, images were acquired on a Zeiss LSM 880 confocal microscope with an Airyscan Fast detector using an EC Plan-Neofluar 40x/1.30 DIC m27 objective. scFv-NG was excited by the 488 nm laser line at 1.2%. Using 2032×1788 pixels and 0.8x optical zoom, 3D optical z-stacks were acquired with 500 nm z-spacing. 55-60 planes were captured with a z-stack acquisition time of ∼ 60 s. The same settings were used for the scFv-NG-KDEL control embryos. Embryos were imaged for 30 min through nc14.

### Quantitation of the number of total and translated *SunTag-hb* mRNAs

For the analysis of zygotic *SunTag-hb* mRNA and translation site numbers, fixed embryos were stained with SunTag smFISH probes and DAPI and imaged using the acquisition details described above. Downstream analysis was carried out in Imaris software (Imaris software 10.1.0 Bitplane, Oxford Instruments) and custom spot assignment scripts as described (Vinter et al., 2021a; Vinter et al., 2021b). Briefly, cytoplasmic mRNAs, scFv-FP/anti-GCN4 signals and translation sites (based on their co-localisation) were identified in 3D by masking nuclear signals, then all signals were assigned to the nearest nucleus. For the quantitation, the proportion of translated mRNAs was calculated for cells with at least 10 (for 24x SunTag) or 5 (for 12xSunTag) mRNAs. Graphs were plotted for data points with at least 1 translation site/nuclear bin.

For the analysis of maternal *SunTag-hb* mRNA and translation site numbers, the analysis was performed as described above for zygotic except that, rather than assigning to the nearest nucleus, the total signals were reported in the region of interest. One of the embryos with 1 copy of the maternal *24xSunTag-hb* transgene had ∼1700 *24xSunTag-hb* mRNAs in the posterior region of interest (and ∼1200 in the anterior), whereas all the other biological replicate embryos with a single copy in Figs. 1 and S1 had <600 mRNAs. Therefore, we excluded the embryo with ∼1700 *24xSunTag-hb* mRNAs from the analysis.

### Ribosome numbers

Scripts for the following analysis are available online from: https://github.com/Hilary-Ashe-Lab/SunTag_Pizzey_2025. To calculate ribosome numbers, the colocalisation of mRNAs and scFv-sfGFP foci was assessed to identify active sites of translation as outlined previously (Vinter et al., 2021a; Vinter et al., 2021b). For colocalised signals, both the sfGFP and mRNA foci were normalised to the intensity of a single molecule (protein or mRNA, respectively) to account for the possibility of multiple proteins and mRNAs existing in a translation site. The length of the SunTag sequence and *hb* CDS are then used in a correction factor to account for ribosomes that had only translated part of the SunTag (Pichon et al., 2016; Vinter et al., 2021a) and the number of ribosomes per mRNA is calculated using the following expression:

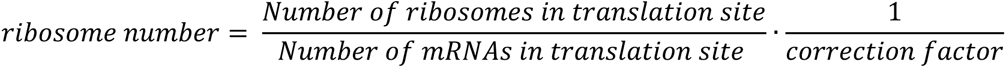

Translation sites were called based on having at least 2 ribosomes.

### Analysis of *SunTag-sog* mRNA translation

Scripts for the following analysis are available online from: https://github.com/Hilary-Ashe-Lab/SunTag_Pizzey_2025. Fixed embryos were stained with SunTag smFISH probes and DAPI as above. Acquired images were analysed in Imaris. *SunTag-sog* mRNA puncta were identified as outlined previously (Vinter et al., 2021a; Vinter et al., 2021b). For the NG foci, due to clustering and the variability in sizes of clusters and individual puncta, spots of volume 0.8 μm in diameter and 1.6 μm in the z direction were placed over the entirety of the embryo to capture as much NG signal as possible. Spot colocalisation, nuclei identification and removal of nuclear spots was performed as outlined (Vinter et al., 2021b). Following removal of nuclear spots, the total SunTag smFISH and total colocalised scFv-NG fluorescence relative to the closest nucleus was calculated across the expression domain. Both SunTag and NG foci were background corrected by fitting spots to background signal in Imaris and subtracting the median intensity of a background spot from all foci. Using data from the central 60% of the embryo, the nuclei were then binned in 12 bins across the DV axis and the average SunTag smFISH and scFv-NG fluorescence per nuclear territory was calculated for each bin. The mRNA per cell was calculated by dividing the intensity of each SunTag foci by the mean intensity of a single SunTag mRNA. The total number of mRNA in all foci was then summed and divided by the number of nuclei in the image. To determine the percentage of translated mRNAs, the total number of SunTag signals colocalised with NG was divided by the total number of SunTag mRNAs in the image.

Live embryos were imaged as described above and movies were processed using Zeiss Airyscan processing. NG intensity levels in the neuroectoderm region were exported from Imaris for each time point. Data sets were aligned with t=0 at the point where nuclei movements had ceased following embryo division from nc13 to nc14.

## Statistical analysis

Statistical comparisons were performed using two-tailed paired or unpaired Student’s *t*-test in GraphPad Prism (Version 10.1.1). Statistical tests used and sample size are described in figure legends, with statistical significance being *P*<0.05.

## Acknowledgements

We thank Martin Pool and Lauren Forbes Beadle for helpful discussions, Daisy Vinter for making the pMatE>24xSunTag-hb plasmid, Sophie Frampton for preliminary imaging of *SunTag-sog* embryos, Mounia Lagha for the scFv-msGFP2 flies, and Golgi Graphics for the figure schematics. We thank Sanjai Patel at the University of Manchester Fly Facility for generating the transgenic flies and the University of Manchester Bioimaging Facility for support.

## Author contributions

Conceptualisation: A.P., M.P.A., H.L.A.; Software: J.C.L.; Investigation: A.P., C.S., E.A.; Writing - original draft: H.L.A., M.P.A., A.P.; Writing - review & editing: A.P., C.S., J.C.L., E.A., M.R., M.P.A., H.L.A.; Supervision: H.L.A., M.R., M.P.A.; Funding acquisition: H.L.A., M.R. M.P.A.

## Data availability

Scripts for image analysis are available from: https://github.com/Hilary-Ashe-Lab/SunTag_Pizzey_2025

