## Supplementary Figures for "Exploiting the SunTag system to study the developmental regulation of mRNA translation"

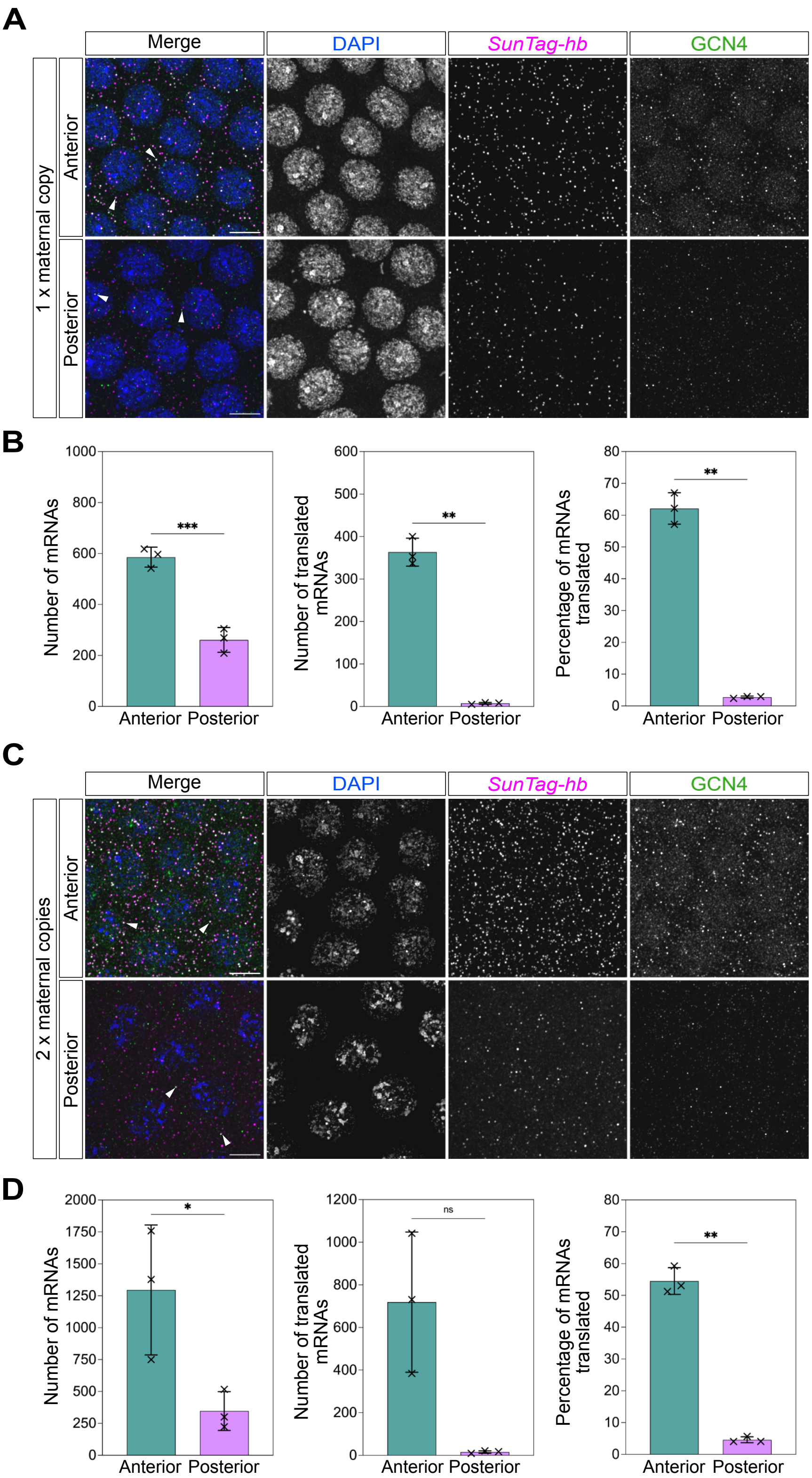

**Fig. S1. Maternal *24xSunTag-hb* mRNA translation in the embryo.**

(A) Representative high magnification images from anterior and posterior regions (as shown in Fig. 1Bi) of a nc13 *matE>24xSunTag-hb*/+ embryo, stained with anti-GCN4 antibody (green), DAPI (blue) and *SunTag* smFISH probes (magenta). White arrowheads show translation sites based on colocalised *24xSunTag-hb* and anti-GCN4 signals. Scale bar: 5 µm.

(B) Quantitation of the number of cytoplasmic mRNAs, translated mRNAs and percentage of translated mRNAs in anterior (cyan) and posterior (magenta) regions of interest in nc13 *matE>24xSunTag-hb*/+ embryos, for 3 biological repeats. Means ± SD, Paired two-tailed Student's *t*-test, ** $P<0.01$, *** $P<0.001$.

(C) As in (A) except images are of a *matE>24xSunTag-hb* homozygous nc13 embryo.

(D) As in (B) for *matE>24xSunTag-hb* homozygous nc13 embryos. ns - not significant, * $P<0.05$, ** $P<0.01$.

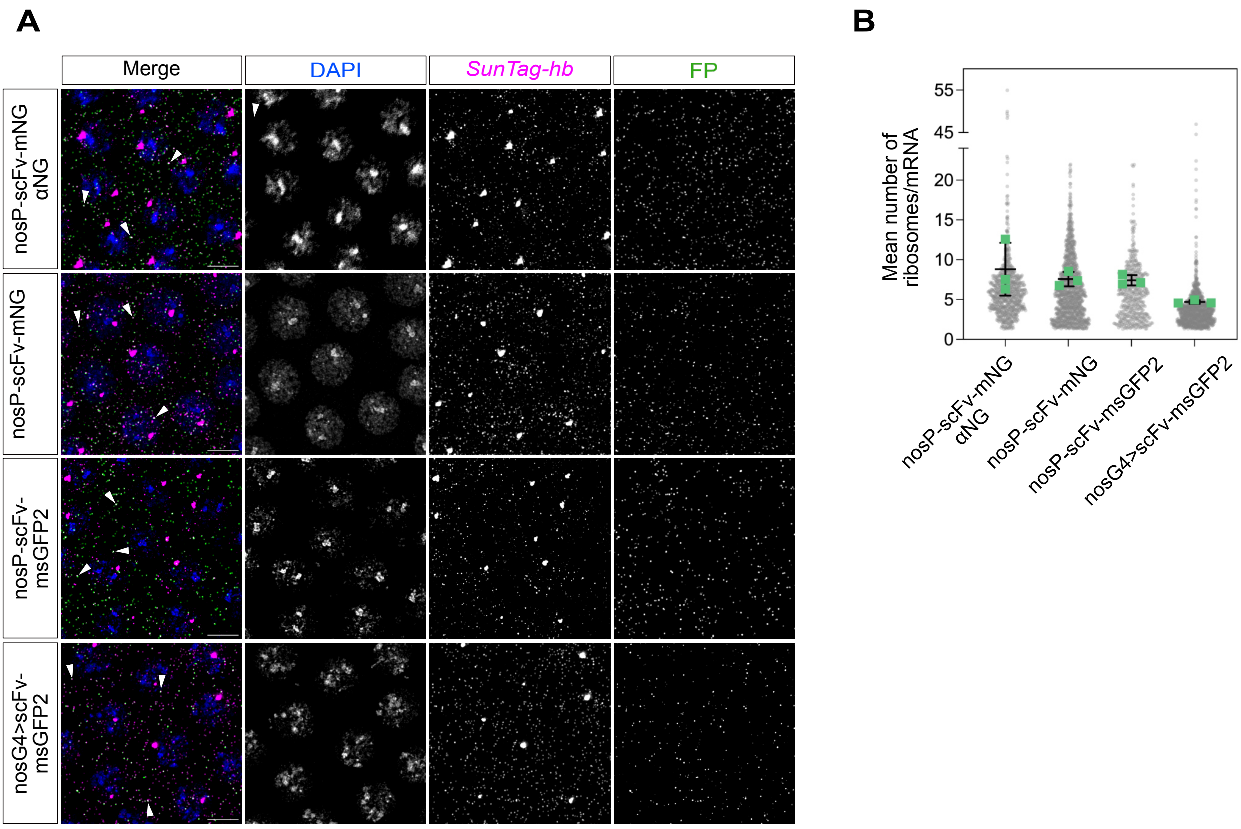

**Fig. S2. Ribosome number analysis in nc13 embryos.**

(A) High magnification images of the anterior of nc13 fixed embryos (region as in Fig. 2Bi) with zygotic *24xSunTag-hb* mRNAs detected by SunTag smFISH probes and translation signals detected as indicated below and summarised in the boxes on the left. Top row: embryos were collected from females expressing *scFv-NG-NLS* from the *nos* enhancer/promoter (nosP) crossed to *hbP2>24xSunTag-hb* males, translation sites were detected by anti-NG antibody staining. Second row: as for the top row, but translation sites were detected by NG fluorescence. Third row: embryos were collected from females expressing *scFv-msGFP2-NLS* from the *nos* enhancer/promoter crossed to *hbP2>24xSunTag-hb* males, translation sites were detected by msGFP2 fluorescence. Fourth row: embryos were collected from *nos-GAL4-VP16*/*UASp-scFv-msGFP2-NLS* females (nosG4>scFv-msGFP2) crossed to *hbP2>24xSunTag-hb* males, translation sites were detected by msGFP2 fluorescence. White arrowheads show translation sites based on colocalised *24xSunTag-hb* and fluorescent protein signals. Scale bar: 5 µm.

(B) Graph shows individual data points and the mean numbers, from 3 biological replicate embryos, of ribosomes present on zygotic *24xSunTag-hb* mRNAs from the embryos described in (A).

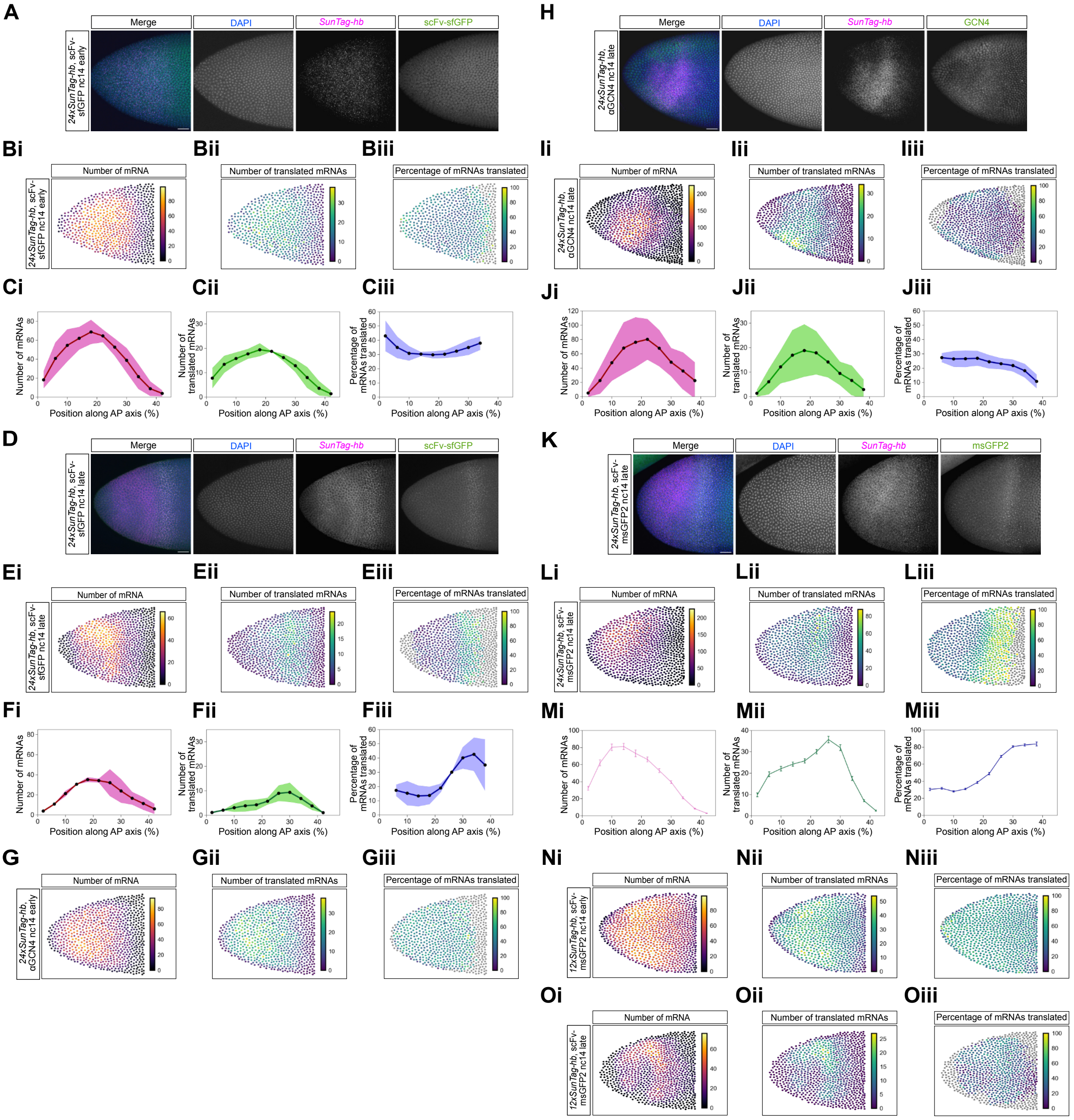

**Fig. S3. Zygotic *24xSunTag-hb* mRNA translation in the embryo.**

(A) Image shows the zygotic *24xSunTag-hb* expression domain in an early nc14 embryo, collected from *matα4-GAL4-VP16 UASp-scFv-sfGFP-NLS* females crossed to *hbP2>24xSunTag-hb* males, showing sfGFP signals (green), and stained with DAPI (blue) and *SunTag* smFISH probes (magenta). Scale bar: 25 µm.

(B) Heatmaps show quantitation, per nuclear territory, of the total mRNAs (i), translated mRNAs (ii) and percentage mRNAs translated (iii) for the embryo shown in (A).

(C) Quantitation of (i) total number of mRNAs, (ii) translated mRNAs and (iii) the percentage of mRNAs translated, per binned nuclear territory, for 3 biological repeats, including the embryo shown in (A). Nuclear territories represent 20 μm bins along the AP axis. Data shown are mean for each bin ± s.d.

(D-F) As in (A-C) except the data are from late nc14 *24xSunTag-hb* embryos.

(G) As in (B) except that the heatmaps correspond to the early nc14 embryo stained with anti-GCN4 shown in Fig. 3B.

(H-J) As in (A-C) except the data are for late nc14 *hbP2>24xSunTag-hb*/+ embryos, stained with anti-GCN4 (green), DAPI (blue) and *SunTag* smFISH probes (magenta).

(K-M) As in (A-C) except the data are for a late nc14 embryo collected from *nos-GAL4-VP16*/*UASp-scFv-msGFP2-NLS* females crossed to *hbP2>24xSunTag-hb* males, showing msGFP2 signals (green), and stained with DAPI (blue) and *SunTag* smFISH probes (magenta). In (M) the data are mean for each bin + S.E.M.

(N, O) As in (B) except that the heatmaps correspond to early (N) and late (O) nc14 *12xSunTag-hb* embryos shown in Figs. 3E and 3G.

(H-J) As in (A-C) except the data are for late nc14 *hbP2>24xSunTag-hb*/+ embryos, stained with anti-GCN4 (green), DAPI (blue) and *SunTag* smFISH probes (magenta).

(K-M) As in (A-C) except the data and graphs are for a late nc14 embryo collected from *nos-GAL4-VP16*/*UASp-scFv-msGFP2-NLS* females crossed to *hbP2>24xSunTag-hb* males, showing msGFP2 signals (green), and stained with DAPI (blue) and *SunTag* smFISH probes (magenta). In (M) the data are mean for each bin + S.E.M from the embryo shown in (K).

(N, O) As in (B) except that the heatmaps correspond to early (N) and late (O) nc14 *12xSunTag-hb* embryos shown in Figs. 3E and 3G.

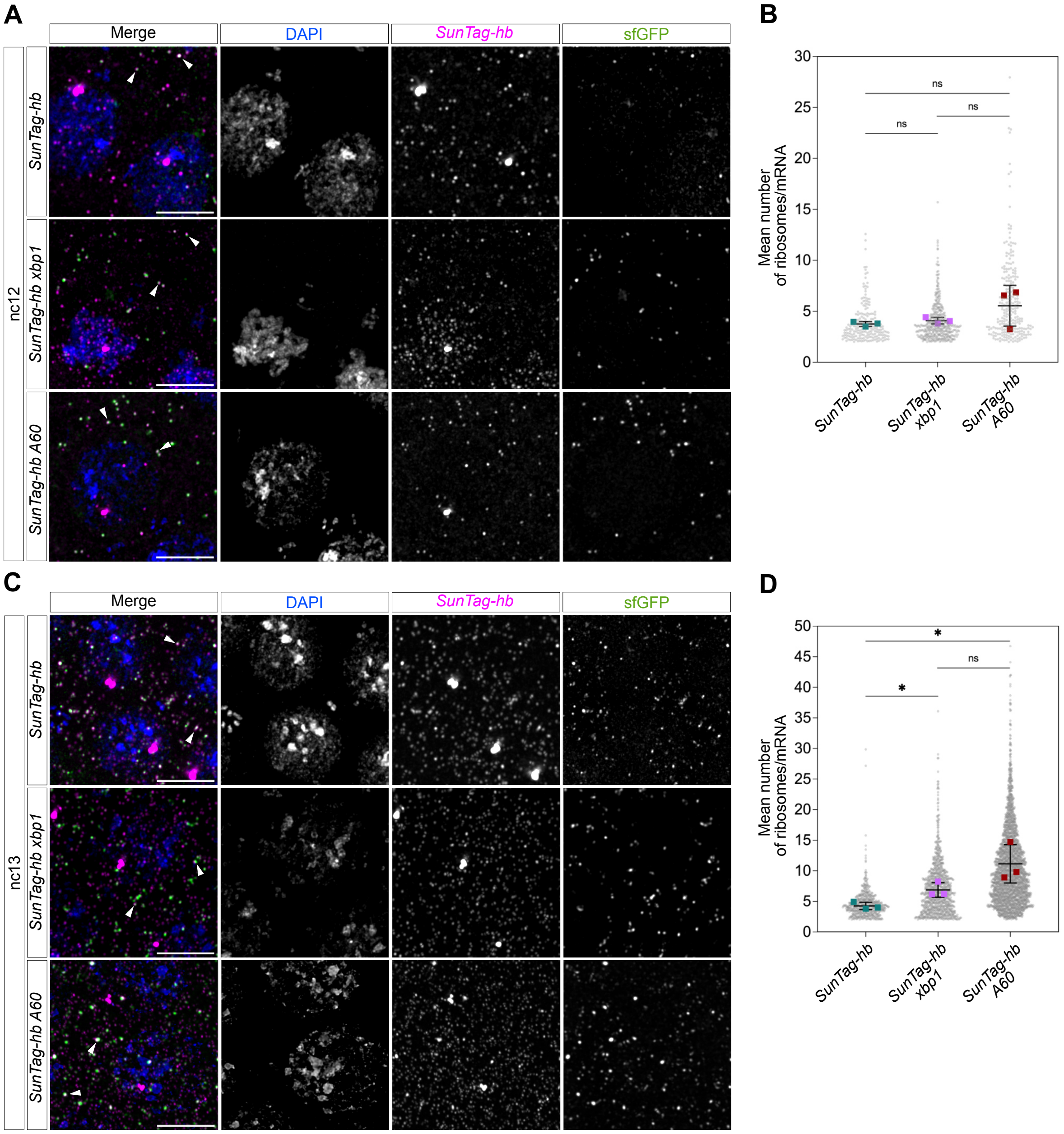

**Fig. S4. Ribosome pausing at nc12 and nc13.**

(A) Representative high magnification images of the anterior region of nc12 embryos collected from *matα4-GAL4-VP16 scFv-sfGFP-NLS* females crossed to *hbP2>24xSunTag-hb*, *hbP2>24xSunTag-hb-xbp1* or *hbP2>24xSunTag-hb-A60* males. Images show sfGFP signals (green), *SunTag* smFISH staining (magenta) and DAPI (blue). White arrowheads indicate translation sites. Scale bar: 5 μm.

(B) Graph shows individual data points and the mean numbers, from 3 biological replicate nc12 embryos, of ribosomes present on zygotic *hbP2>24xSunTag-hb*, *hbP2>24xSunTag-hb-xbp1* or *hbP2>24xSunTag-hb-A60* mRNAs, unpaired two-tailed *t*-test, ns not significant, **P*<0.05, ***P*<0.01. Data are mean ± s.d.

(C-D) As in (A, B) except the data are for nc13 embryos.

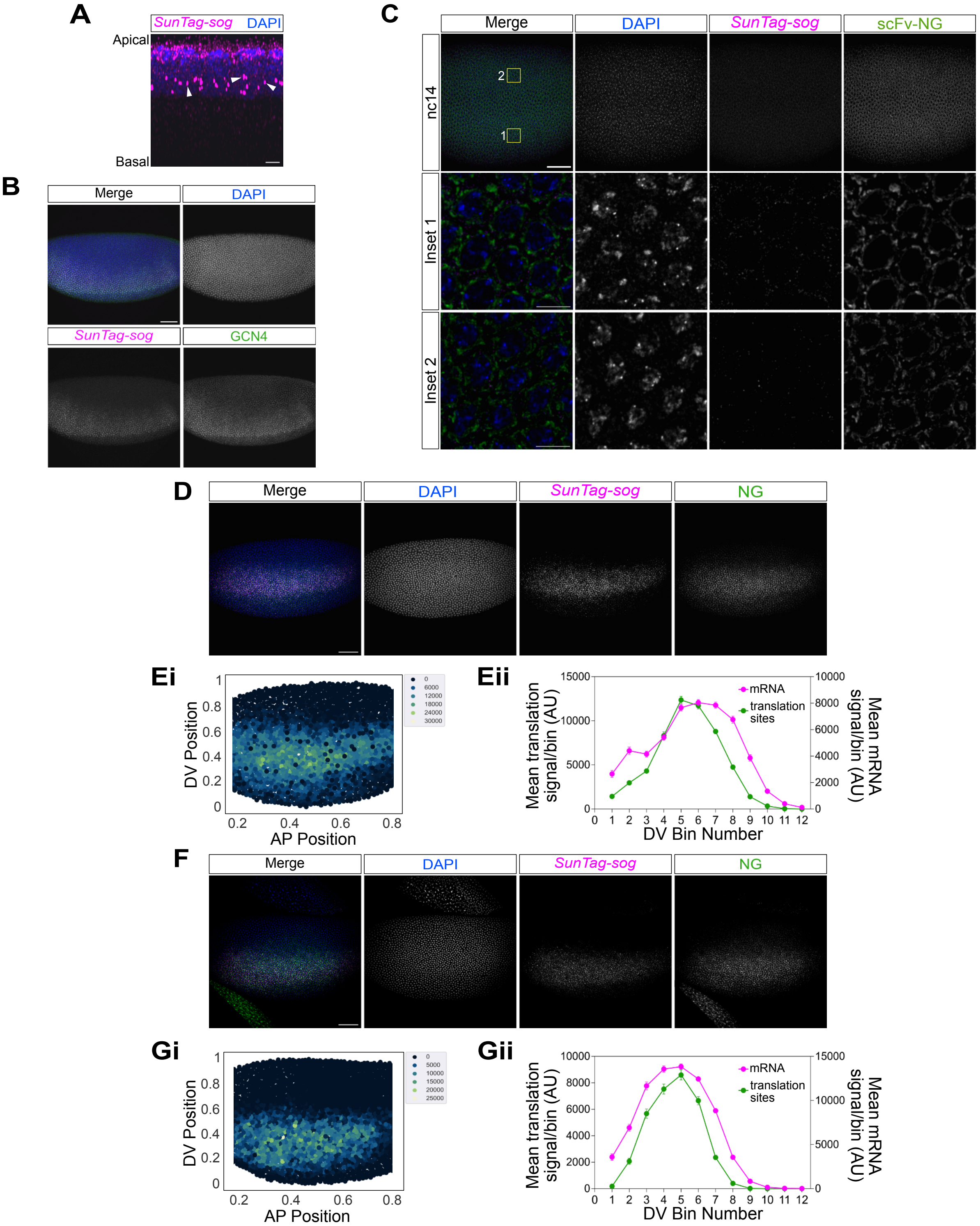

**Fig S5. Visualising translation of SunTag-sog mRNA translation in fixed embryos.**

(A) An orthogonal view of a nc14 *SunTag-sog* embryo stained with SunTag smFISH probes (magenta) and DAPI (blue). The *SunTag-sog* mRNAs are apically localised in the cell, transcription sites (white arrowheads) are located basally in the nuclei. Scale bar: 2 μm.

(B) Image of a nc14 *SunTag-sog* embryo stained with SunTag smFISH probes (magenta), anti-GCN4 (green) and DAPI (blue). Scale bar: 50 μm.

(C) Central region of a fixed nc14 embryo from females carrying *nos-GAL4-VP16* and *UASp-BIP-scFv-NG-KDEL* insertions crossed to control males, showing NG signals (green) and stained with DAPI (blue) and *SunTag* smFISH (magenta). The embryo regions used for high magnification images are shown. Scale bar: 50 μm for lateral view, 5 μm for insets.

(D, F) Lateral views of additional biological repeat nc14 embryos, from females carrying *nos-GAL4-VP16* and *UASp-BIP-scFv-NG-KDEL* crossed to *SunTag-sog* males. Embryos are stained with DAPI (blue) and *SunTag* smFISH probes (magenta), with NG signals (green). Scale bars: 50 μm.

(E, G) Heatmaps showing the total NG signal per cell (i), and quantitation of the mean NG and mRNA signals per binned cell (ii). Cells are represented in 12 bins across the DV axis and the graphs show the mean value of each bin ± S.E.M.

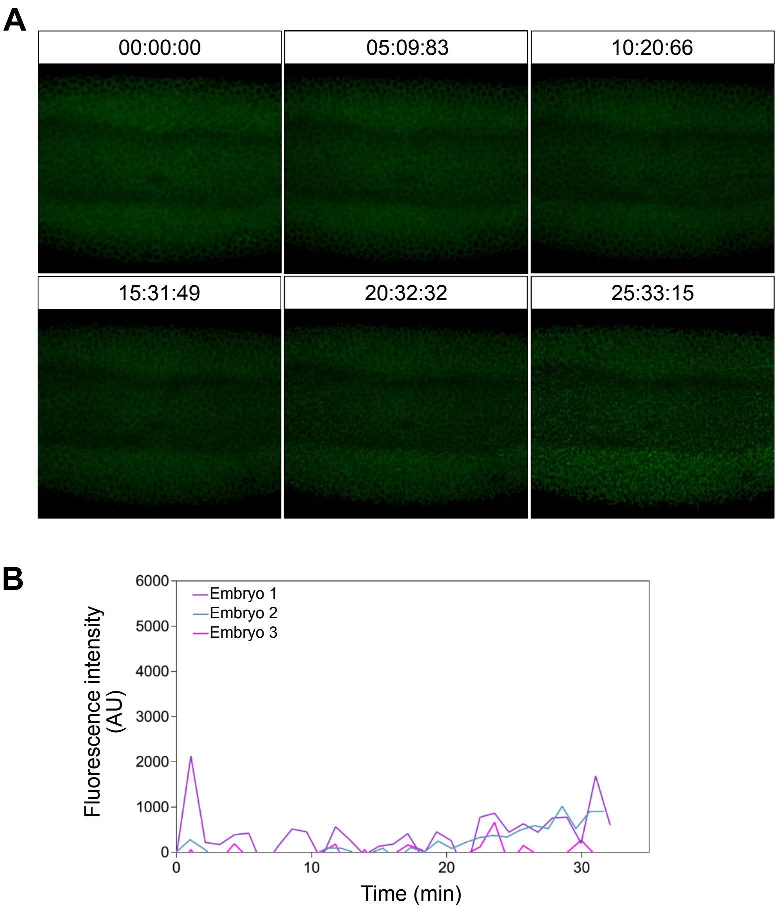

**Fig S6. Analysis of control *nos-GAL4-VP16*/*UASp-BIP-scFv-NG-KDEL* embryos.**

(A) Stills from a movie of a control embryo, collected from females carrying *nos-GAL4-VP16* and *UASp-BIP-scFv-NG-KDEL* insertions crossed to control males, developing through nc14. Time intervals are min:s:ms. See Movie 2.

(B) Quantitation of NG signals in the neuroectoderm from 3 control embryos over a ~30 min period in nc14.

**Supplementary Movies.**

Movie 1. Movie of a representative embryo (ventrolateral view) showing *SunTag-sog* translation (green). NG signal was quantitated over a ~30 min period in nc14. The embryo was collected from females carrying *nos-GAL4-VP16* and *UASp-BIP-scFv-NG-KDEL* insertions crossed to *SunTag-sog* males. Imaged with a 40x objective, 0.8 optical zoom and ~60 sec time resolution per frame. Scale bar: 30 μm. Time intervals are min:s:ms.

Movie 2. Movie of a representative control embryo, collected from females carrying *nos-GAL4-VP16* and *UASp-BIP-scFv-NG-KDEL* insertions crossed to wildtype males. Imaging settings are as for Movie 1.

**Table S1. Primers used in this study.**

| Primer | Sequence |
|---|---|
| hbP2 StuI F | TCGAGGTCATCAAGCTTAGGCCTTAACTACGAAACTGCCCACGC |
| hb CDS BamHI R | ATGGTGATGGGGAACGGATCCTTAGGAGTGAGCATTCCTGGCC |
| 12SunTag CDS F | CTTAAAGAAACAGAACTGGGAGACGACAGCC |
| 12SunTag R | CCCAGTTCTGTTTCTTTAAGCGCGCGACTTCG |
| Pause hb CDS R | CCTGCATCTTGGAGTGAGCATTCCTGGCC |
| Pause F | TGCTCACTCCAAGATGCAGGCAACGCAGAG |
| Pause BamHI R | ATGGTGATGGGGAACGGATCCTTAGGCCATTAGCTCTATGCCCG |
| p60A R | ATGGTGATGGGGAACGGATCCTTATTTTTTTTTTTTTTTTTTTTTTTTTTTTT<br>TTTTTTTTTTTTTTTTTTTTTTTTTTTTTTTGGAGTGAGCATTCCTGGCC |
| Hb CDS NotI F | TAGACTCGAGCGGCCGCCAGAACTGGGAGACGACA |
| Hb 3’UTR R | AGCACAGTGGCGGCCGAATATGTTAGGTTAAATAAGAC |
| Exon 1 NotI Oligo F | CCGGCCGTCTAGAGCCGCCAAGATGGC |
| Exon 1 NotI Oligo R | GGCCGCCATCTTGGCGGCTCTAGACGG |
| Exon 2 F | TGTCCGCAAGCCGTCTAGAGCCGCCAAGA |
| Exon 2 NdeI R | TCAGCTCCAAAGCACGCATACGAATATGTTAGGTTAAATAAGACT |
| MatE HindIII F | TTTCTCGAGGTCATCAAGCTCCTTAAGGAGATTTGGAATTTCGA |
| MatE R | CTCTAGACGGCTTGCGGACAGTCCAAGTG |
| SunTag NotI F | GGGGCGGCCGCGGAAGAACTTTTGAGCAAG |
| SunTag NotI R | GGGGGCGGCCGCCCTTTTTAAGTCGGGCTACTTC |
| mNeonGreen F | AGGCGGCGGAAGCTTGGATCCAGGTGGAGGTGGAAGCGG |
| GB1 R | CTTAGTACTTCTAGAGGATCCTTATCAAAGCTCGTCCTTACTAGTACCACC<br>ACCGCTACC |
| BiP ScFV F | CGGGGATCAGATCCGCGGCCGCTCTCAATATGAAGTTATGCATATTACTGG<br>CCGTCGTGGCCTTTGTTGGCCTCTCGCTCGGGATGGGCCCCGACATCGT<br>G |
| Tubulin 3’UTR R | ACGTTCGAGGTCGACTCTAGAAAATGACATCAGACATAACCTCAA |
| (A/B) LS EcoRI F | GGGCGCGTACTCCACGAATTCTGGTCCATGGTCCATACC |
| (A) 5’UTR mid R | CACACAATGGATATAGATATATCAGGTGTA |
| (A) 5’UTR mid F | ATATCTATATCCATTGTGTGTGCCAGTGTG |
| (A) Sog TM R | CACCACCACCCATGAGCGGCGCATGCCG |
| (A) GCN4 F | GCCGCTCATGGGTGGTGGTGGAAGTGGC |
| (A) GCN4 EcoRI R | TGCGGCCGCTCCGGAGAATTCCTTTTTAAGTCGGGCTACTTCATTC |
| (B) GCN4 R | CACCACCACCCTTTTTAAGTCGGGCTACTTCATTC |
| (B) Sog CDS after TM F | ACTTAAAAAGGGTGGTGGTGGAAGTGGTG |
| Sog R | TGGAGCCGCCTTAGCTGGAGGATCGCTGCTG |
| Sog 3’UTR F | CTCCAGCTAAGGCGGCTCCACGTGACGG |
| (B) Sog 3’UTR EcoRI R | TGCGGCCGCTCCGGAGAATTCATGGGTATATTTCGAATATATTTTG |
